# Engineering Adaptive Alleles for *Escherichia coli* Growth on Sucrose Using the EasyGuide CRISPR System

**DOI:** 10.1101/2024.12.20.629655

**Authors:** Joneclei Alves Barreto, Matheus Victor Maso Lacôrte e Silva, Danieli Canaver Marin, Michel Brienzo, Ana Paula Jacobus, Jonas Contiero, Jeferson Gross

## Abstract

Adaptive Laboratory Evolution (ALE) is a powerful approach for mining genetic data to engineer industrial microorganisms. This evolution-informed design requires robust genetic tools to incorporate the discovered alleles into target strains. Here, we introduce the EasyGuide CRISPR, a five-plasmid platform that exploits *E. coli*’s natural recombination system to assemble gRNA plasmids from overlapping PCR fragments. The production of gRNAs and donor DNA is further facilitated by using recombination cassettes generated through PCR with 40 to 60-mer oligos. With the new CRISPR toolkit, we constructed 22 gene edits in *E. coli* DH5α, most of which corresponded to alleles mapped in *E. coli* DH5α and E2348/69 ALE populations selected for sucrose propagation. For DH5α ALE, sucrose consumption was supported by the *cscBKA* operon expression from a high-copy plasmid. During ALE, plasmid integration into the chromosome, or its copy number reduction due to the *pcnB* deletion, conferred a 30–35% fitness gain, as demonstrated by CRISPR-engineered strains. A ∼5% advantage was also associated with a ∼40.4 kb deletion involving *fli* operons for flagella assembly. In E2348/69 ALE, inactivation of the *hfl* system suggested selection pressures for maintaining λ-prophage dormancy (lysogeny). We further enhanced our CRISPR toolkit using yeast for in vivo assembly of donors and expression cassettes, enabling the establishment of polyhydroxybutyrate synthesis from sucrose. Overall, our study highlights the importance of combining ALE with streamlined CRISPR-mediated allele editing to advance microbial production using cost-effective carbon sources.

## 1. Introduction

Adaptive Laboratory Evolution (ALE) is a dynamic area of biological research where organisms undergo evolutionary changes in controlled laboratory environments (Barrick and Lenski, 2013). By selecting for adaptive mutations and enhancing phenotypes, ALE serves as a powerful tool for developing industrial microbial strains with increased production capacity and improved stress tolerance (Dragosits and Mattanovich, 2013; Menegon et al., 2022). Moreover, the application of next-generation sequencing to ALE populations or evolved clones facilitates the precise identification of adaptive alleles that are enriched under selection. This knowledge allows for reverse-engineering evolved mutations back into parent strains, providing insights into allele-specific fitness and guiding the rational design of industrial strains (Barrick and Lenski, 2013; Menegon et al., 2022).

*Escherichia coli* is a preferred microorganism for biotechnological applications due to its ease of cultivation and robust tools for genetic modification (Blount, 2015; Jiang et al., 2015). However, most *E. coli* strains are unable to utilize sucrose, a cost-effective carbon source derived from abundant feedstocks like sugarcane and beet crops (Bruschi et al., 2012). To enable sucrose consumption, industrial and laboratory strains can be engineered with the *cscB*, *cscK*, and *cscA* genes, which are naturally found in the operons of *E. coli* W and enteropathogenic strains (Bruschi et al., 2012; Olavarria et al., 2019). However, implementing a heterologous pathway often requires ALE to ensure seamless integration into the host’s metabolic network and to identify genetic targets that can be reverse-engineered to enhance sucrose utilization (Mohamed et al., 2019).

Reconstructing ALE mutations necessitates the use of robust genetic tools (Menegon et al., 2022). Over the past decade, CRISPR/Cas9 has emerged as the premier genome editing technology, enabling gene deletions, the integration of heterologous pathways, and precise single-nucleotide modifications (Dong et al., 2021). Numerous studies have highlighted strategies for genome editing in *E. coli* using CRISPR/Cas9 (Table 1). These typically involve plasmid-based co-expression of *Streptococcus pyogenes* Cas9 and the λ-Red Exo/Bet/Gam recombinase system (Dong et al., 2021). This phage-derived recombinase enhances the chromosomal integration of donor DNAs in *E. coli*, serving as templates for genome edits (Bassalo et al., 2016; Jiang et al., 2015).

**Table 1.** Key CRISPR approaches using Cas9 endonuclease for genome edition in *E. coli*.

| Reference | Cas9 plasmid and gRNA cloning | Donor production |
| --- | --- | --- |
| (Jiang et al., 2013) | Two-plasmid system (Crispr RNA and Cas9).<br>gRNA cloned via restriction-ligation. | Linear oligonucleotide donor. |
| (Jiang et al., 2015) | pTargetF (gRNA) and pCas9 (Cas9). gRNA cloned<br>by inverse PCR, or restriction-ligation. | Plasmid donor (250/550 bp HA), cloned by overlap<br>PCR and restriction-ligation. Linear PCR Donors. |
| (Li et al., 2021) | Two-plasmid system. gRNA complementary oligos<br>ligated into Bsal-prepared plasmid. | Linear donor. |
| (Li et al., 2023) | Two-plasmids. p15A-Cas9 (high-copy number).<br>gRNA oligos ligated into a Bsal-cut plasmid. | Linear donor. |
| (Reisch and Prather, 2015) | Two-plasmid system (gRNA/λ-Red and Cas9).<br>gRNA cloned via CPEC. | Donors as oligonucleotides: 60 – 90 nts ssDNA or<br>dsDNA oligos. |
| (Pyne et al., 2015) | Three-plasmid system (gRNA, Cas9, and λ-Red).<br>gRNA oligos ligated into Bsal-prepared plasmid. | 60-nts ssDNA donors. dsDNA donors from *lacZ*<br>amplification by PCR. The primers provide 40 bp HA. |
| (Bassalo et al., 2016) | gRNA plasmid assembled through CPEC. Four plasmids for gRNA, Cas9, λ-Red, and donor. | Circular donors cloned with CPEC (50 – 600 bp HA). Linear 10 kb integration cassette cloned in yeast. |
| (Zerbini et al., 2017) | Two-plasmid system. gRNA oligonucleotides ligated into a Bsal-prepared plasmid. | ssDNA and ds DNA oligos (70 nt and 120 nts). |
| (Shukal et al., 2022) | gRNA prepared through restriction-free cloning. Two- plasmid system (gRNA and Cas9/ λ-Red). | Linear donors with asymmetric HA (50 bp / 500 bp) prepared by PCR. |
| (Zhao et al., 2016) | gRNA plus donor assembled via Golden Gate. One plasmid for Cas9, λ-Red, gRNA and donor. | Donor with 41 – 300 bp HA cloned with the gRNA via Golden Gate assembly. |
| (Li et al., 2015) | Two-plasmid system. PCR-amplified plasmid with primer-encoded gRNA, Golden Gate assembly. | Donors with 300 – 500 bp HA produced via fusion PCR. |
| (Fang et al., 2024) | Combined plasmid for gRNA and Cas9. gRNA cloning through MetClo assembly. | Plasmid donor cloned through MetClo assembly. |
| (Wang et al., 2021) | Two-plasmid system. gRNA cloned via PCR. Rock-Paper-Scissors approach for plasmid curing. | Donor cloned into the gRNA plasmid by using seamless clone kits or restriction-ligation. |
| (Lim et al., 2023) | Cas9-harboring strain. gRNA plasmid cloning via Gibson assembly. Use of 5'-truncated gRNAs. | Donors as 41–70-mer oligonucleotides. |
| (Feng et al., 2018) | Three-plasmid system (Cas9/λ-Red, gRNA, and donor). Golden Gate assembly. | Donors cloned and delivered within plasmids. Golden Gate assembly. |
| (Moreb et al., 2017) | Two-plasmid system (gRNA/λ-Red and Cas9/recA56). gRNA cloning with commercial kits. | Donors as ssDNA and dsDNA oligos. |
| (Sun et al., 2019) | Two-plasmid system (Cas9/λ-Red and gRNA). gRNA cloning through inverse PCR. | Linear donors assembled through fusion PCR. |
| (Tang et al., 2017) | Three-plasmid system (Cas9, λ-Red and gRNA/SceI). gRNA cloned through PCR. | Linear dsDNA donor supplied as a PCR product assembled through overlap extension PCR. |
| (Zhao and Guo, 2024) | Two plasmids: Cas9/Beta and gRNA, cloned by restriction-ligation and homologous recombination. | Linear dsDNA generated through extension overlap PCR. |

However, recent research indicates that RecA-proficient *E. coli* strains can achieve high-efficiency gene editing with attenuated Cas9/Cas12 targeting (Collias et al., 2023). A crucial element of the CRISPR protocol is the engineering of guide RNAs (gRNAs), which harbor a 20-nt spacer that directs Cas9 to its genomic targets (Jacobus et al., 2022). Each planned genome modification requires the cloning of a new gRNA into a separate plasmid, which is then introduced into a host strain expressing the Cas9/λ-Red recombinase system (Table 1).

The construction of donor DNAs is also a major bottleneck in CRISPR/Cas9 experiments, primarily due to the need for assembling large homology arms (HA) of 100- 600 bp flanking the editing sequence (Bassalo et al., 2016; Jiang et al., 2015). This often requires precloning, increasing both cost and time of the genome edit. Despite numerous studies outlining various strategies for cloning gRNAs and donors (Table 1), there is still a demand for simple, cost-effective techniques (Shukal et al., 2022). In vivo cloning in *E. coli* provides a practical and economical alternative to traditional restriction-ligation methods, potentially eliminating the need for expensive cloning kits based on Gibson and Golden Gate assemblies (Watson and Garcia-Nafria, 2019). This method leverages *E. coli*’s natural ability to recombine DNA fragments with short overlapping homologies (10- 50 bp) (Jacobus and Gross, 2015; Kostylev et al., 2015). Although the precise mechanisms of this gap-repair cloning are not fully understood, it likely involves a RecA-independent pathway with 3’ to 5’ exonucleolytic activity (ExoIII) generating single-stranded DNA (ssDNA) termini (Conley et al., 1986; Nozaki and Niki, 2019). These ssDNA ends can anneal and be repaired, possibly by DNA polymerase I (*polA*) and DNA ligase (Nozaki and Niki, 2019).

In this study, we conducted an ALE experiment to improve *E. coli* growth using sucrose as the sole carbon source. To streamline the CRISPR/Cas9 workflow for reverse-engineering ALE mutations, we adapted the EasyGuide approach—originally developed for yeast—for *E. coli* (Jacobus et al., 2022). We introduce a five-plasmid platform to facilitate PCR generation of DNA components for assembling gRNA-encoding plasmids via in vivo cloning in *E. coli*. Additionally, we present the EasyOligo approach, enabling straightforward PCR amplification of gRNAs and donors. Our strategies for in vivo donor assembly in yeast also facilitates the construction of heterologous expression cassettes, advancing the metabolic engineering of polyhydroxybutyrate (PHB) production from sucrose in *E. coli*.

## 2. Materials and methods

### 2.1 Strains and growth conditions

*E. coli* DH5α, E2348/69, and yeast strains are listed in Supplementary Table 1. The purity of *E. coli* cultures was confirmed using MacConkey agar (Merck KGaA, Darmstadt, Germany) and Gram staining (Laborclin, Pinhais, PR, Brazil). *E. coli* was cultured in liquid LB medium (1% tryptone, 0.5% yeast extract, 1% NaCl) at 37 °C with 200 rpm shaking or on solid medium with 1.5% agar. Antibiotics used included ampicillin (100 µg/mL), kanamycin (50 µg/mL), spectinomycin (50 µg/mL), and chloramphenicol (50 µg/mL). Cells were stored in 25% glycerol at -80 °C. Yeast cells were cultured in YPS medium (1% yeast extract, 2% peptone, 2% sucrose) with hygromycin (300 µg/mL) or nourseothricin (100 µg/mL) as needed.

### 2.2 PCR and molecular cloning procedures

PCR oligonucleotides were ordered from Exxtend Biotechnology (Campinas, Brazil) and are listed in Supplementary Tables 2 and 3. DNA fragments for cloning and transformations were produced by PCR with Phusion High-Fidelity DNA Polymerase (Thermo Fisher Scientific, Waltham, MA, USA) or Genie Fusion Ultra High-Fidelity DNA Polymerase (Assay Genie, Dublin, Ireland). GoTaq™ DNA Polymerase (Promega, Madison, WI, USA) and Platus Taq DNA Polymerase (Sinapse Inc, São Paulo, Brazil) were used for diagnostic PCRs. PCR products were analyzed on 1–3% agarose gels in TBE buffer with 0.01% ethidium bromide. For colony PCR, cells from a colony were heated in water, and the lysate was used for diagnostic PCRs. PCR fragments were purified with the QIAquick PCR Purification Kit or QIAquick Gel Extraction Kit (QIAGEN, Hilden, Germany) as necessary.

In the EasyOligo method for generating gRNAs and donor DNAs, oligos were prepared at 10 pmol/µL. For PCR, 2.5 µL of flanking oligos and 1.25 µL of the central oligo were used. PCR involved an initial denaturation at 95 °C, 30 cycles of 95 °C for 10 seconds, 50 °C for 15 seconds, and 72 °C for 10 seconds, with a final extension at 72 °C for 1 minute. Products were analyzed via electrophoresis on 3% agarose gels.

Cloning in *E. coli* DH5α employed the in vivo technique (Jacobus and Gross, 2015). For each fragment, 5 µL of the PCR reaction was used for transformation. If needed, 10 U of DpnI (New England Biolabs, Ipswich, MA, USA) was added to the PCR tube.

Competent cells were prepared using the rubidium chloride method (Hanahan, 1983). For heat shock transformations (Maniatis et al., 1982), DNA-mixed cells were incubated at 42 °C for 1 minute, chilled on ice for 20 minutes, then recovered in 1 mL LB medium at 37 °C with shaking at 200 rpm for 1 hour. After recovery, they were centrifuged, plated on LB agar with antibiotic, and incubated at 37 °C for 12 to 16 hours. For *E. coli* electroporation (Lessard, 2013), 500 µL of overnight culture was grown in 50 mL LB until OD600 reached 0.5-0.7. Cells were centrifuged, washed twice with 10% glycerol, resuspended in 500 µL of 10% glycerol, and aliquoted into 50 µL portions for electroporation or freezing. Electroporation was performed at 2.5 kV for 5 ms in a MicroPulser electroporator (Bio-Rad, Hercules, California, USA) using 0.2 cm cuvettes. Post-electroporation, cells were recovered in 1 mL LB at 37 °C with shaking for 1 hour, and 50 µL was plated on antibiotic media for selection.

Plasmids are listed in Supplementary Table 4. For DNA plasmid preparation, *E. coli* DH5α cultures were grown in 10 mL of LB medium with the appropriate antibiotic at 200 rpm for 12–16 hours. Plasmids were extracted using the QIAprep® Miniprep kit (QIAGEN, Hilden, Germany) and quantified with a Qubit 2.0 Fluorometer (Life Technologies, Carlsbad, CA, USA). Plasmid DNA was stored at -20 °C.

For cloning donor DNA plasmids in yeast, the 2µ replication region and resistance markers were amplified from plasmids pEasyG3-mic, pEasyG3-hph, and pEasyG3-nat (Jacobus et al., 2022) using the primer pairs 2Mic_f/2micMX and tMX_2m/MXProUni, respectively. Transformations used 5 µL of PCR products introduced into *Saccharomyces cerevisiae* CEN.PK113-7D via the lithium acetate method (Gietz and Woods, 2002); transformed cells were selected on YPS plates with hygromycin or nourseothricin. Plasmids were enriched from *S. cerevisiae* using a QIAprep® Miniprep kit (QIAGEN, Hilden, Germany) after yeast lysis with zirconia beads.

### 2.3 Genomic modifications through the EasyGuide CRISPR system

Detailed procedures for constructing plasmids pTargetAmp, pSpec5’, pSpec3’, pOliSpec5’, and pOliSpec3’ are provided in Supplementary File 1. These plasmids are available at Addgene (https://www.addgene.org/) under the catalog numbers 228548– 228552, and their maps are displayed in Supplementary Figs. 1–5.

To assemble gRNA plasmids using pTargetAmp as a template, PCR was performed with primers P1 and P2, both containing 20-nt spacers. Plasmid pSpec5’ was amplified with P2/P3, and pSpec3’ with P1/P4. PCR products for pTargetAmp, pSpec5’, and pSpec3’ were transformed into *E. coli* DH5α via heat shock (Maniatis et al., 1982). For pOliSpec5’ and pOliSpec3’, PCR used P2/P3 and P1/P4, respectively, with P1 and P2 serving as priming regions only. These products were cotransformed with oligo-amplified gRNAs, and successful cloning was confirmed by diagnostic PCR with primers A and B. The gRNA oligonucleotides are listed in Supplementary Table 3, with gRNA assembly details in Supplementary File 1.

The temperature-sensitive plasmid CAS9BAC1P-1EA (Merck KGaA, Darmstadt, Germany), encoding the *Streptococcus pyogenes* Cas9 and λ-Red recombinase, was transformed into *E. coli* DH5α and used according to the manufacturer’s instructions. An overnight DH5α culture with CAS9BAC1P-1EA was diluted 1:100 in 50 mL LB containing kanamycin and grown at 30 °C and 200 rpm. Upon reaching an OD600 of 0.2–0.3, L-arabinose was added to 10 mM. Cells at OD600 0.5–0.7 were centrifuged and resuspended in 2 mL. Aliquots of 50 µL were used for the heat shock transformation (Hanahan, 1983), or occasional use of electroporation (Lessard, 2013).

For CRISPR/Cas9 experiments, 50 µL cells with the CAS9BAC1P-1EA plasmid were transformed via heat shock with 10 µL of donor PCR product and 40 ng of gRNA plasmid. Controls included only the gRNA plasmid and the pSpec plasmid. After recovery in LB with 10 mM L-arabinose at 30 °C and 200 rpm, cells were plated on LB agar containing kanamycin, spectinomycin, and L-arabinose. Colony PCRs confirmed genetic modifications. The gRNA plasmid was removed through successive LB passages without antibiotics, and the CAS9BAC1P-1EA was eliminated by propagation at 37 °C. All genetic modifications are described in Supplementary File 1.

### 2.4 Adaptive laboratory evolution

*E. coli* E2348/69 (Iguchi et al., 2009) was transformed with pKan, pSpec, pAmp, and pGGA plasmids (Supplementary File 1) to create four distinct ALE populations: E-Kan, E- Spec, E-Amp, and E-Cm. *E. coli* DH5α was first transformed with the pCscBKA plasmid (Supplementary Fig. 6), followed by the pKan, pSpec, and pGGA plasmids to produce the D-Kan, D-Spec, and D-Cm populations.

ALE was conducted by serially propagating each population in minimal medium with 2% sucrose (Supplementary Methods). The process included daily 1:100 dilutions in 20 mL of medium within 50 mL Erlenmeyer flasks at 37 °C with shaking at 80 rpm. OD600 was measured daily. Each population was maintained on its respective antibiotic to prevent contamination, and ampicillin was used for DH5α populations to maintain pCscBKA. Purity was periodically checked using MacConkey agar and Gram staining. Weekly glycerol stocks were made, and propagation lasted about three months.

### 2.5 Genome DNA sequencing and variant calling

Genomic DNA from the final and progenitor populations was purified using the phenol:chloroform (1:1) extraction (Maniatis et al., 1982). DNA quality was checked with 1% agarose gel electrophoresis, and quantification was done with a Qubit 2.0 Fluorometer (Life Technologies, Carlsbad, CA, USA). Libraries were prepared with the NEBNext® Ultra™ II DNA Library Prep Kit (New England BioLabs, Ipswich, MA, USA), and 2 x 150 bp paired-end sequencing (Q30 > 85%) was performed on the Illumina NovaSeq 6000 platform (GenOne, Solutions in Biotechnology, Rio de Janeiro, Brazil). Sequencing reads were deposited in NCBI under BioProject number PRJNA1200122.

Reads were filtered to retain those with a Phred quality score of at least 30 and a length of 75 bases. Mapping was conducted using the Burrows-Wheeler aligner in CLC Genomics Workbench 8.01 (QIAGEN, Aarhus, Denmark) with a cutoff of 0.8 for both read length and identity (Jacobus et al., 2024). DH5α reads were mapped to the DH5α reference genome (GenBank CP017100), and E2348/69 reads were mapped to their reference chromosomes (GenBank FM180568, FM180570, FM180569), excluding duplicate reads. Variants with a frequency of ≥ 30% specific to evolved populations were identified. Read mappings against plasmids were also performed. Plasmid copy numbers for each ALE population were calculated by normalizing read coverage over antibiotic resistance markers to their length and dividing by genomic read coverage (Jacobus et al., 2021).

### 2.6 Phenotypic analyses

Growth curves of parent and evolved populations were measured using a Tecan Sunrise microplate reader (Tecan, Männedorf, Switzerland). After two preadaptation passages in minimal medium with 2% sucrose, cells were counted with an Attune NxT flow cytometer (Thermo Fisher Scientific, Waltham, MA, USA). A 5 µL dilution (∼10^4^ cells) was used to inoculate 200 µL of the same medium in a 96-well microplate, with four replicates per population. Growth at 37 °C was monitored by recording OD600 every 15 minutes. Maximum specific growth rates (µmax) were determined during the exponential phase by plotting the natural logarithm of OD600 against time and fitting a linear regression. The µmax for each replicate was determined from the slope and averaged (Rodrigues et al., 2021).

To assess the fitness of evolved DH5α populations, D-Kan, D-Cm, and D-Spec strains (expressing pCscBKA with AmpR) were competed against a tester-GFP strain (*SS9:cscBKA; lacZ*::*gfp; galK::gfp*) with respective marker plasmids plus pAmp, which conferred ampicillin resistance. Initial tests showed that the DH5α ALE progenitor (carrying the pCscBKA) was quickly outcompeted by the tester-GFP. To address this, after preadaptation passages, we mixed the tester-GFP and the ALE progenitor at a 95%:5% ratio, while evolved populations were mixed in a 1:1 ratio with the tester-GFP. From these mixtures, we inoculated 200 µL into 20 mL of 2% sucrose minimal medium in triplicate using 125 mL Erlenmeyer flasks. During two growth passages (1:100 dilutions) at 37 °C and 100 rpm, GFP-expressing and non-expressing cells were measured at the stationary phase using an Attune NxT flow cytometer.

The cell proportions allowed us to calculate the fitness coefficient (S) using the formula: S = ln[ALEf/GFPf] - ln[ALEi/GFPi] (Jacobus et al., 2024). Here, ALEf/GFPf are the final ALE populations and tester-GFP cell proportions, while ALEi/GFPi are the initial proportions of the competitors after the first passage. Cell counts at each point allowed us to normalize the fitness per cell doubling (S/doubling). The S/doubling obtained for the progenitor DH5α-pCscBKA competing against tester-GFP was used to calculate the fitness burden from pCscBKA expression and to normalize the S/doubling of evolved ALE populations. The cost of GFP expression was also calculated in a competition between the DH5α *SS9::cscBKA* strain and tester-GFP.

Evolved E2348/69 populations were also competed against the DH5α tester-GFP. However, as evolved E2348/69 populations exhibited much faster growth rates, the competitions were set up with a 10:1 proportion of the tester-GFP to evolved E2348/69, while the E2348/69 progenitor was competed starting from a 1:1 mixture. Competitions and S/Doubling calculations were performed under the same conditions as described above.

Fitness evaluation for genetically modified strains started from a 1:1 strain/tester- GFP mixture and was recorded during four passages, with the first one serving as a baseline for S/doubling calculations. Normalization was obtained with the S/doubling calculated from a competition of *SS9::cscBKA* versus the tester-GFP. Statistical analyses were performed using GraphPad Prism 8.01 (GraphPad Software, La Jolla, California, United States).

We measured the fluorescence of the pGFP plasmid transformed into DH5α, *ΔpcnB*, and D-Spec backgrounds by recording 10,000 events with the Attune NxT flow cytometer after triplicate overnight cultures and IPTG induction. Fluorescence intensity values were derived from the median of flow cytometry histograms and averaged (Jacobus et al., 2024).

Procedures for sucrose and PHB quantification are detailed in the Supplementary Methods.

## 3. Results

### 3.1 ALE experiment

Many *E. coli* strains that naturally utilize sucrose carry the *cscB*, *cscK*, and *cscA* genes within an operon (Fig. 1A). The *cscB* gene encodes a sucrose/H^+^ symporter for the transport of the disaccharide into the cell, where CscA invertase hydrolyzes it into fructose and glucose. CscK kinase phosphorylates fructose to fructose-6-phosphate, a glycolytic intermediate (Bruschi et al., 2012; Olavarria et al., 2019). The *E. coli* strain E2348/69 (Iguchi et al., 2009) contains the *cscBKA* operon. We used *E. coli* in vivo recombination (Jacobus and Gross, 2015) to clone this operon into the pUC backbone, creating the pCscBKA plasmid (Fig. 1A and Supplementary Fig. 6). The functionality of pCscBKA in DH5α was confirmed by its capacity to support growth in minimal medium supplemented with sucrose (Supplementary Fig. 7).

**Fig. 1.**
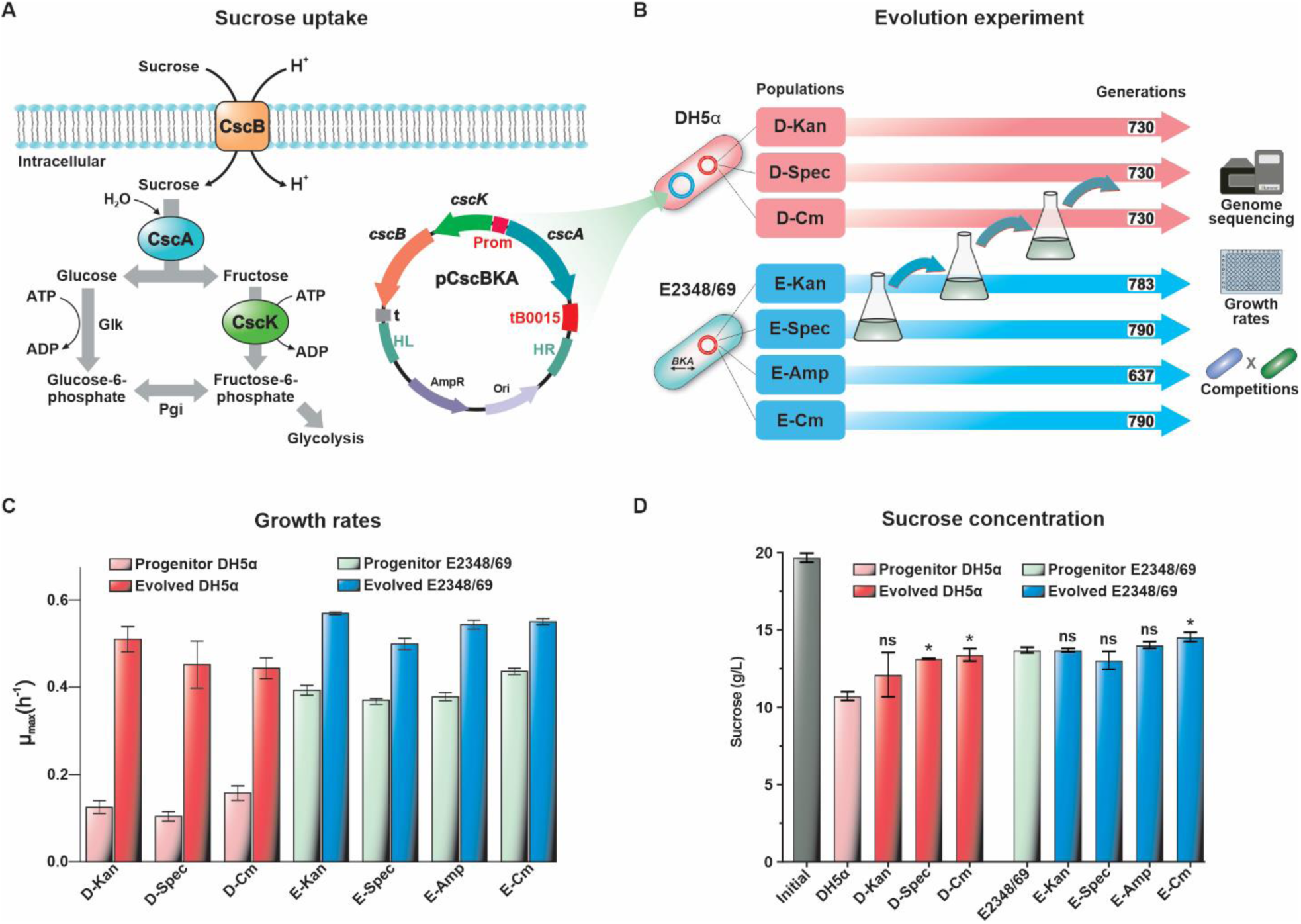
ALE experiment in 2% sucrose. (A) Sucrose catabolic pathway in DH5α with the pCscBKA plasmid: *cscB*, *cscA*, and *cscK* encode a sucrose transporter, invertase, and fructokinase, respectively. (B) ALE setup: DH5α and E2348/69 populations, tagged with plasmid resistance to Kan, Spec, Amp, or Cm, evolved via daily 1:100 transfers on 2% sucrose for 637– 790 generations. Final populations underwent DNA sequencing and growth/competition assays. (C) µmax comparisons for progenitor versus evolved populations in 96-well microplates. (D) Sucrose consumption analysis with an initial concentration of 20 g/L (grey bar) and the remaining sucrose concentration in the medium after 24 h of culture. The DH5α progenitor was transformed with the pCscBKA plasmid. Statistical significance: ns (non-significant); (*) *p* < 0.05.

To enhance the growth of DH5α containing pCscBKA on sucrose, we developed an ALE protocol propagating *E. coli* populations on this sugar for hundreds of generations over approximately 120 days (Fig. 2A). Three distinct DH5α ALE populations were tagged with unique antibiotic resistance plasmids: D-Kan, D-Spec, and D-Cm. To explore alternative evolutionary pathways for improved sucrose utilization, we also established four E2348/69 ALE populations, each distinguished by a plasmid-encoded antibiotic resistance marker: E-Kan, E-Spec, E-Cm, and E-Amp. These antibiotic resistance genes and their corresponding antibiotics also provided protection against contamination during the ALE process.

**Fig. 2.**
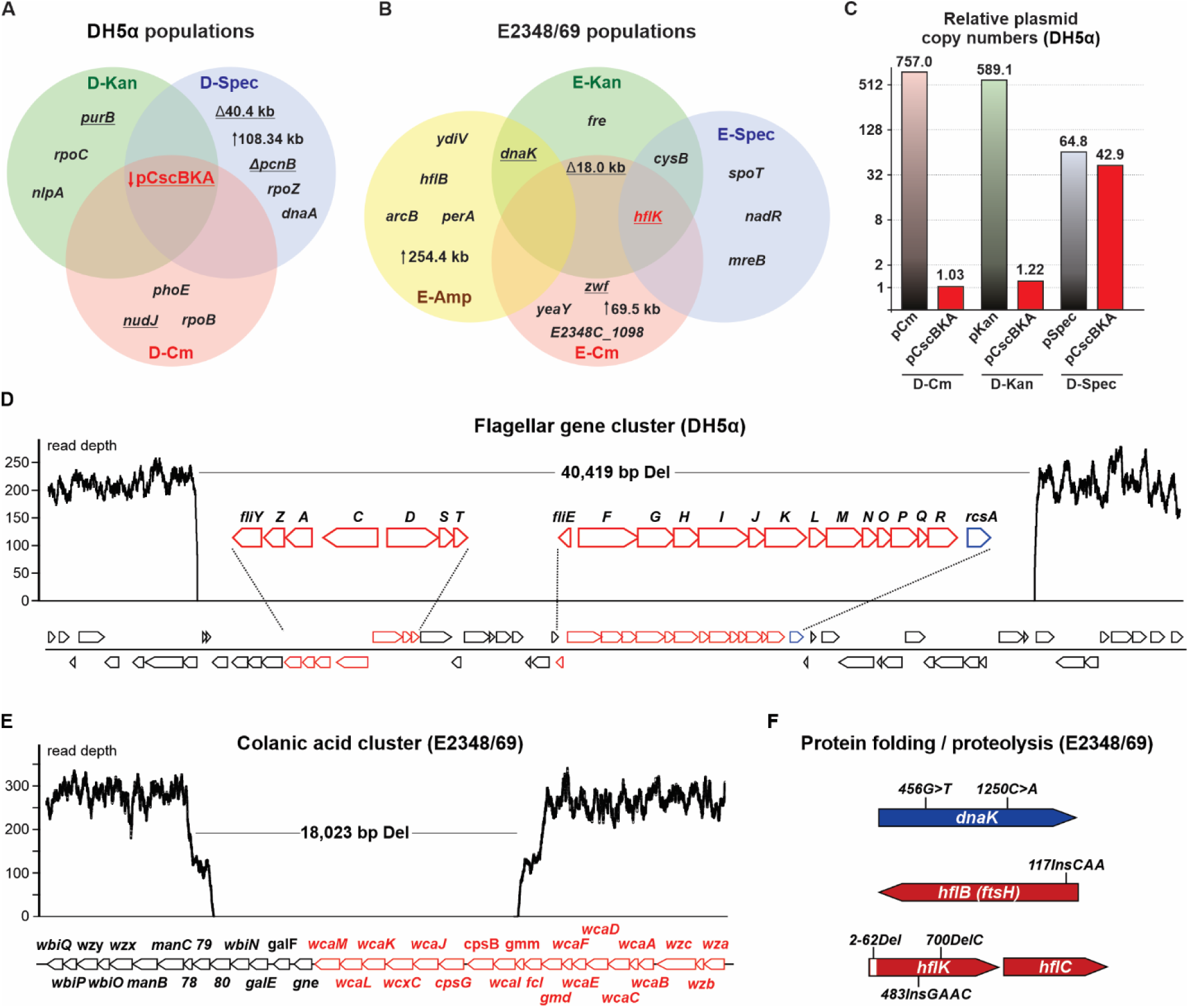
Summary of mutations in ALE populations. (A, B) Venn diagrams illustrating common and strain-specific mutations; upward arrows indicate segmental amplifications, while downward arrows signify reductions in copy number. Underlined alleles have been selected for reverse engineering. The most frequent targets of the evolution are highlighted in red. (C) Illumina read depth analysis estimating plasmid copy numbers relative to the chromosome in final DH5α ALE populations. (D) Deletion of a ∼40.4 kb segment in the DH5α D-Spec population, encompassing flagellum assembly clusters (*fliA-R* and *fliYZ* genes, red). The *rcsA* gene (blue) is a transcription regulator for CA synthesis. (E) An 18,023 bp deletion in E-Kan and E-Cm, affecting part of the CA biosynthetic cluster (red). (F) Mutations in genes related to protein folding and proteolysis: *dnaK* (E-Kan, E-Amp), *hflB* (E-Amp), and *hflK* (E-Kan, E-Spec, E-Cm).

Upon completion of the experiment, genomic DNA from all final populations was sequenced using the Illumina platform, and phenotypes were analyzed (Fig. 1C and D; Supplementary Fig. 7). The maximum specific growth rate (µmax) of the evolved populations increased compared to their progenitors, with a more pronounced increase in the DH5α populations (Fig. 1C). However, the DH5α populations exhibited decreased total sucrose consumption relative to the parent, while the E2348/69 populations showed no significant change in final sucrose concentration compared to the progenitor. These findings suggest that enhanced sucrose consumption was not the primary driver of evolution.

### 3.2 Genome Sequencing of Evolved Populations

DNA sequencing of the evolved populations and their progenitors was conducted on the Illumina NovaSeq 6000 platform, identifying mutations such as SNPs, segmental amplifications/deletions, and small InDels (summarized in Fig. 2A and B, and Supplementary Table 5). Notably, Illumina read depth for the pCscBKA sequence in DH5α D-Kan and D-Cm populations was similar to that of the *E. coli* chromosome (Fig. 2C), unlike typical high-copy pUC plasmids that replicate hundreds of copies per cell (Lee et al., 2006). This suggests potential integration of pCscBKA into the chromosome during ALE. Examination of Illumina read mappings revealed broken read pairs over the pCscBKA sequence in D-Kan and D-Cm, with unaligned ends linking to insertion sequences IS1 (D-Kan) and IS4 (D-Cm), indicating possible genomic integrations mediated by these mobile elements (Supplementary Fig. 8).

A different pattern was observed in the evolved D-Spec population, where the coverage depth over the pCscBKA and pSpec plasmids decreased by ∼10-fold compared to pKan and pCm (Fig. 2C). Additionally, an 81 bp deletion in *pcnB* was detected, a mutation known to reduce the copy number of ColE1 plasmids (Lopilato et al., 1986), such as the pCscBKA.

Further functionally overlapping mutations in DH5α populations involved RNA polymerase subunits RpoC, RpoB, and RpoZ, which are linked to a trade-off of increased fitness in minimal media but reduced growth in rich media (Conrad et al., 2010; Mohamed et al., 2019). The D-Spec population had a ∼40.4 kb deletion affecting flagellar assembly gene clusters (Fig. 2D). In D-Kan, a SNP in *purB* caused an L115G change, potentially repairing a mutation known to lower DH5α growth rates in minimal medium (Jung et al., 2010). Additional mutations in DH5α populations were found in the genes *dnaA*, *phoE*, and *nudJ*.

The evolved alleles in E2348/69 populations are summarized in Fig. 2B. An identical ∼18 kb deletion in the colanic acid (CA) biosynthetic cluster was found in both E-Cm and E-Kan populations (Fig. 2E). Similar parallelism appeared in the *hfl* system, with different *hflK* alleles in E-Kan, E-Spec, and E-Cm, and a CAA insertion at position 117 in *hflB* (Fig. 2F). The *hflKC* operon encodes membrane proteins that regulate HflB (FtsH), a protease involved in the turnover of substrates such as the lambda phage CII transcriptional factor, which facilitates phage lysogeny, and the σ^32^ protein, which manages the heat shock response (Bandyopadhyay et al., 2011; Kihara et al., 1996; Tomoyasu et al., 1995).

Parallelism was observed for *dnaK* mutations (E-Kan and E-Amp, Fig. 2F), encoding a chaperone that serves as a protein stability quality control hub (Calloni et al., 2012). SNPs were identified in the *cysB* promoter region in E-Kan and E-Spec populations, which regulate cysteine synthesis (Lilic et al., 2003). Mutations in *zwf* and *nadR* were detected in E-Cm and E-Spec populations, respectively. The *zwf* gene encodes glucose-6- phosphate dehydrogenase, involved in the oxidative branch of the pentose phosphate pathway to generate NADPH (Zhao et al., 2004). NadR functions as a regulator of NAD^+^ biosynthesis and in the uptake of nicotinamide mononucleotide (NMN), the precursor of NAD^+^ (Raffaelli et al., 1999).

### 3.3 Development of a CRISPR/Cas9 strategy based on the EasyGuide principle

To better understand the adaptive role of key ALE mutations, we established a CRISPR/Cas9 platform for the rapid introduction of evolved alleles into parental strains. Genomic modifications via CRISPR/Cas9 were performed using a two-plasmid system with linear DNA donors. The temperature-sensitive plasmid CAS9BAC1P (Sigma-Aldrich), with optimal replication at 30 °C, expressed *S. pyogenes* Cas9 and the λ-Red recombinase. Specific strategies were developed for the production of gRNAs and donor DNAs.

We previously detailed the EasyGuide approach for the in vivo assembly of functional gRNAs into a vector separate from the Cas9 plasmid (Jacobus et al., 2022). This method uses homologous recombination of PCR products in *S. cerevisiae*, eliminating the need for in vitro ligation. We have now adapted the EasyGuide principle for *E. coli*, enabling in vivo gRNA assembly through three different cloning strategies with varying plasmid sets (Figs. 3A-C).

**Fig. 3.**
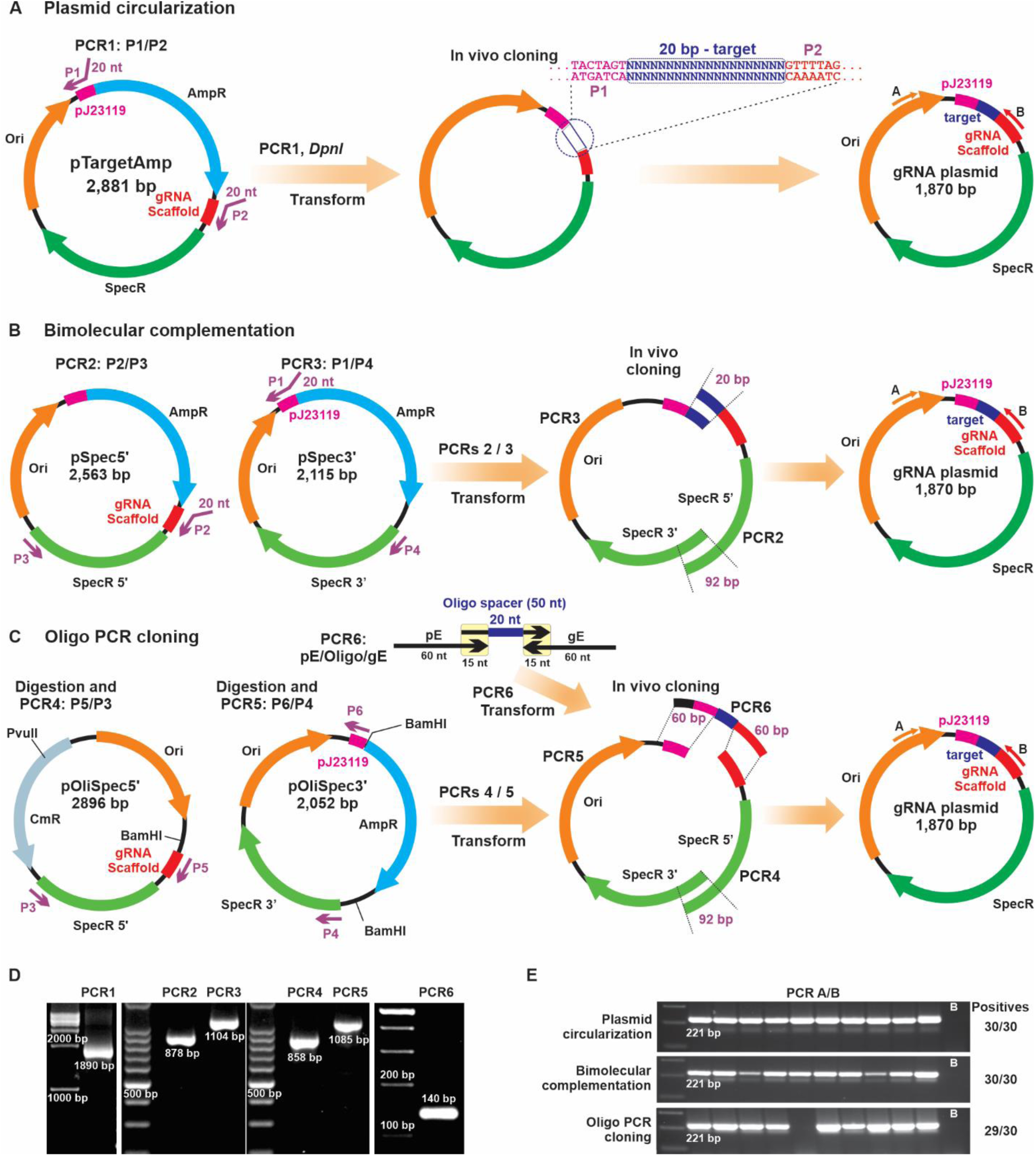
Different strategies for producing gRNA plasmids. (A) Plasmid circularization: using pTargetAmp as a template, PCR is performed with primers P1 and P2 (PCR1, 1,890 bp), specifying the 20-nt spacer. After DpnI treatment, the product is transformed and ligated in vivo. (B) Bimolecular complementation: PCR fragments from pSpec5’ (primers P2/P3) and pSpec3’ (primers P1/P4) are cotransformed into *E. coli*, assembling the gRNA plasmid in vivo. (C) Oligo PCR cloning: plasmids pOliSpec5’ and pOliSpec3’ are digested and purified, then amplified by PCR using primers P5/P3 and P4/P6. The gRNA region is generated by amplifying the oligo spacer with primers pE and gE. The 140 bp product is mixed with backbone fragments and transformed into *E. coli* for in vivo assembly. (D) Electrophoresis in 1%, 1.5%, and 3% agarose gels showing PCR products for gRNA plasmid cloning. Sizes are indicated (1 kb marker for PCR1, 100 bp ladder for PCRs 2–6). (E) 2% agarose gel electrophoresis showing 221 bp PCR products confirming *galK* gRNA cloning using the three strategies. Triplicate tests were conducted; only one replicate is shown. Positive cloning rates are indicated; a 100 bp ladder was used as a marker.

The first gRNA assembly strategy involves plasmid circularization (Fig. 3A). The pTargetAmp plasmid, derived from pTargetF (Jiang et al., 2015), maintains the ColE1 origin and spectinomycin resistance marker, with an ampicillin resistance marker added between the prJ23119 promoter and the gRNA scaffold for spacer sequence insertion. PCR amplifies the pTargetAmp backbone using primers P1 and P2, which have 20-nt 5’ overhangs specifying the spacer. After producing an 1,890 bp product (Fig. 3D), DpnI is added to eliminate the template. The linear plasmid is then transformed by heat shock into *E. coli*, where in vivo recombination circularizes it, ligating the spacer. The resulting plasmid expresses a functional gRNA and confers spectinomycin resistance.

Colonies exhibiting ampicillin resistance represent false positives and were excluded from validation, which employs PCR with primers A and B to produce a 221 bp product in positive clones (Fig. 3E). Validated gRNA plasmids are purified and cotransformed with the linear donor into DH5α cells containing the Cas9/λ-Red recombinase plasmid. This system successfully cloned gRNAs for *galK* (Fig. 3E), *purB*, and the SS9 region (Table 2). However, for some gRNAs, cloning attempts failed, and in other cases, a high rate of false positives was observed.

**Table 2.** CRISPR/Cas9-mediated genetic modifications through the EasyGuide approach.

| Genetic modification | gRNA cloning | Donor assembly | Efficiency |
| --- | --- | --- | --- |
| *SS9::cscBKA* | Plasmid circularization | In vivo cloning in *E. coli* | 10/10 |
| *galK::cscBKA* | Plasmid circularization | In vivo cloning *S. cerevisiae* | 10/10 |
| *cscA::sucP* | Bimolecular complementation | In vivo cloning *S. cerevisiae* | 4/4 |
| *cscA::pJ23119-sucP* | Bimolecular complementation | In vivo cloning *S. cerevisiae* | 4/4 |
| *lacZ::sucP* | Bimolecular complementation | In vivo cloning *S. cerevisiae* | 3/14 |
| *galK::gfp* | Plasmid circularization | In vivo cloning *S. cerevisiae* | 10/10 |
| *lacZ::gfp* | Bimolecular complementation | In vivo cloning *S. cerevisiae* | 1/10 |
| *lacZ::phaCAB* | Bimolecular complementation | In vivo cloning *S. cerevisiae* | 6/14 |
| *galK::ampR* | Plasmid circularization | In vivo cloning *S. cerevisiae* | 5/5 |
| *lacZ::ampR* | Bimolecular complementation | In vivo cloning *S. cerevisiae* | 3/5 |
| $\Delta 40.4$ kb | Bimolecular complementation | PCR Amplification from D-Spec DNA | 4/4 |
| $\Delta fliA$ (205 bp) | Bimolecular complementation | In vivo cloning *S. cerevisiae* | 10/10 |
| $\Delta fliA$ (205 bp) | Bimolecular complementation | Oligo PCR amplification | 9/10 |
| *ΔflhDC* | Oligo PCR cloning | Oligo PCR amplification | 10/10 |
| *ΔpcnB* (81 bp) | Bimolecular complementation | PCR amplification from the D-Spec DNA | 1/10 |
| *ΔhflK* | Oligo PCR cloning | PCR of a donor with asymmetric arms | 10/10 |
| *ΔCA* (12 kb) | Oligo PCR cloning | Oligo PCR amplification | 9/10 |
| *ΔgalK* (32 bp) | Oligo PCR cloning | Oligo PCR amplification | 10/10 |
| *nudJ* (G151V) | Bimolecular complementation | In vivo cloning *S. cerevisiae* | 2/2 |
| *purB* (L115G) | Plasmid circularization | In vivo cloning *S. cerevisiae* | 5/5 |
| *dnaK* (T417N) | Oligo PCR cloning | Oligo PCR amplification | 3/10 |
| *zwf* (D405E) | Oligo PCR cloning | Oligo PCR amplification | 9/9 |

To overcome these problems, a second system was designed using a bimolecular complementation strategy (Fig. 2B). In this system, the gRNA plasmid is functional only through the recombination of two PCR products. The coding sequence for *aadA*, which confers spectinomycin resistance (SpecR) (Murphy, 1985), was divided between two plasmids (Fig. 3B). PCR of pSpec5’ with primers P2/P3 (PCR2: 878 bp, Fig. 3D) amplifies the gRNA and *aadA* 5’ region (SpecR 5’), while PCR of pSpec3’ with primers P1/P4 (PCR3: 1,104 bp, Fig. 3D) amplifies the prJ23119 module and *aadA* 3’ end (SpecR 3’). Cotransformation of these PCR products results in in vivo recombination, producing a functional gRNA plasmid that expresses full-length *aadA* (SpecR). This system avoids plasmid carryover, eliminates the need for DpnI treatment, and minimizes false positives. Using this approach, we have achieved 100% efficiency in gRNA cloning attempts.

To streamline the gRNA plasmid cloning procedure, a third strategy based on oligo PCR cloning (the EasyOligo approach, Fig. 3C and D) was developed. This enhances the bimolecular complementation method by enabling the mass preparation and storage of the PCR products (SpecR 5’/gRNA and SpecR 3’/prJ23119) for use as plasmid backbones in multiple gRNA assemblies utilizing PCR products generated from specific oligos.

For this purpose, two new plasmids, pOliSpec5’ and pOliSpec3’, were constructed with restriction sites to remove the plasmid template backbone prior to PCR amplification (Fig. 3C). This digestion prevents improper recombination events during in vivo cloning (Barreto, personal observations). The pOliSpec5’ plasmid is digested with PvuII and BamHI, while the pOliSpec3’ plasmid is digested with BamHI. After purification, these templates are PCR-amplified using primers P3/P5 (pOliSpec5’, PCR4) and P4/P6 (pOliSpec3’, PCR5) (Fig. 3D). Primers P5 and P6 do not include 5’-overhangs; instead, the 20-nt spacer is embedded within a central oligonucleotide amplified with primers pE (for recombining with prJ23119) and gE (overlapping with the gRNA scaffold). The 140 bp dsDNA produced (Fig. 3C and D, PCR6) is transformed with PCR products 4 and 5. Proper recombination of all three fragments assembles a SpecR gRNA plasmid.

### 3.4 Donor DNA assembly in yeast

Previous CRISPR/Cas9 protocols for *E. coli* have recommended using DNA donors with flanks of 100–600 bp for efficient homologous recombination (Bassalo et al., 2016; Jiang et al., 2013). We developed a strategy to preassemble donors with large homologous regions using yeast in vivo cloning (Fig. 4). The yeast origin of replication (2µ) and an antibiotic resistance marker were amplified by PCR from the pEasyG3 plasmid series (Fig. 4A-C) (Jacobus et al., 2022). Homologous flanking regions are PCR-amplified from the chromosomal target site (Fig. 4A), along with the gene or DNA sequence intended for integration. Oligonucleotides used for PCR create 20–40 bp overlaps between products and connections to the yeast plasmid backbone (Fig. 4A). Deletion borders or SNPs are specified between the homologous arms (Figs. 4B and C). PCR products and plasmid fragments are cotransformed into *S. cerevisiae* without DNA cleanup. After selection, positive colonies are verified by PCR. Donor plasmid enrichment occurs through yeast plasmid preparation. PCR amplification from this preparation yields a donor DNA fragment ready for the CRISPR/Cas9 experiment in *E. coli* via heat shock transformation. An example of this approach is the cloning of a GFP cassette into the *galK* locus of *E. coli* DH5α (Fig. 4D-E).

**Fig. 4.**
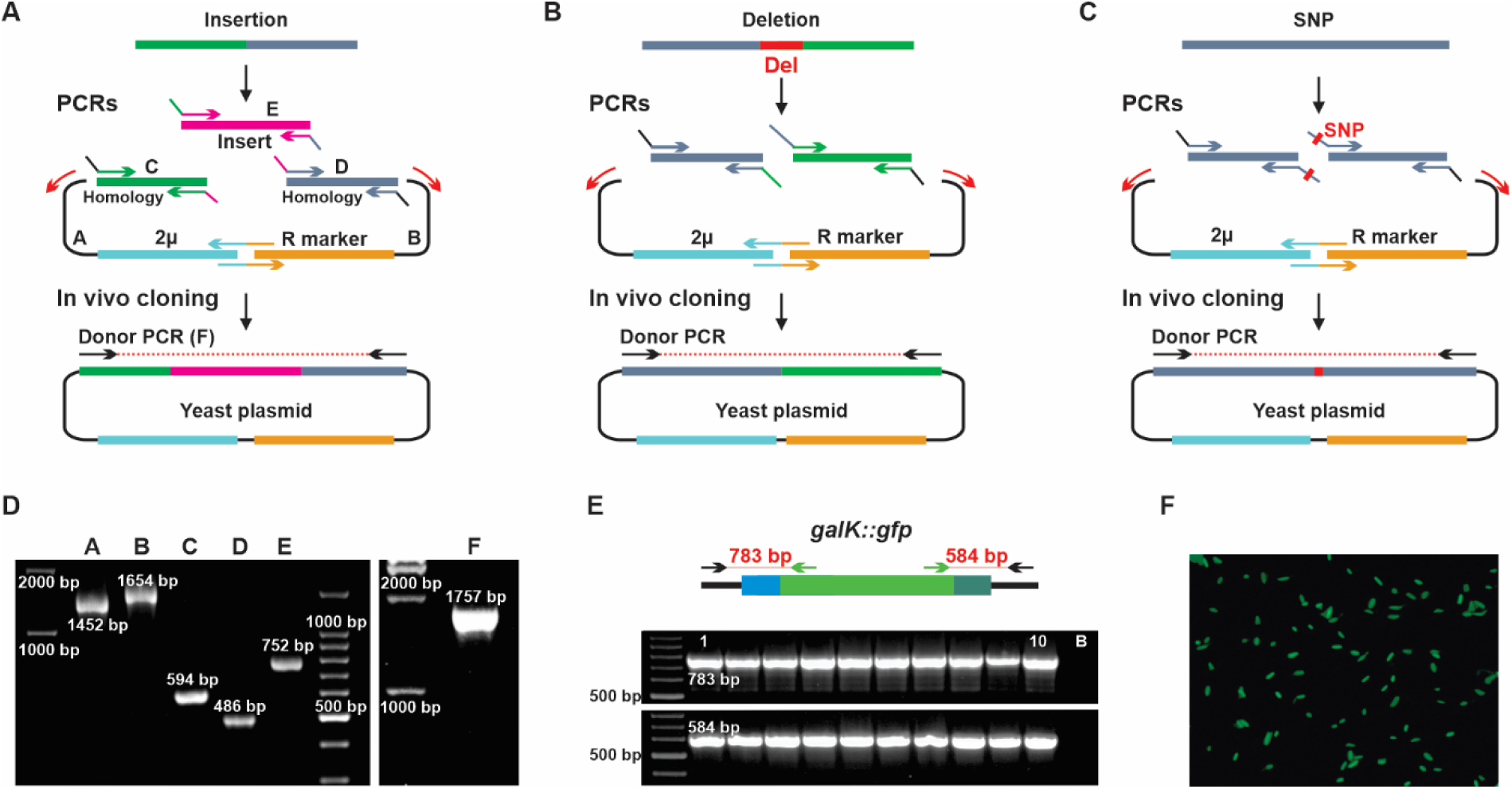
Assembly of donor sequences by yeast homologous recombination. The yeast plasmid origin of replication (2μ) and antibiotic resistance marker (R marker) are amplified from pEasyG3 plasmids via two PCRs (A and B). (A-C) Parts are recombined in vivo with PCR products to create plasmids with donors. (A) Donor DNA precloning for the *E. coli* genome insertion. (B) Donor DNA assembly for chromosomal deletions, positioning the deletion border between homologous arms. (C) Donors for reverse engineering SNPs, with SNPs specified at the 5’ ends of oligonucleotides used for PCR. Assembled donors are PCR-amplified and transformed into *E. coli* for CRISPR/Cas9 integration. (D) A 1.5% agarose gel showing PCRs of parts (A-E) for inserting GFP into the *galK* locus; donor PCR yields 1,767 bp product (PCR F). 1 kb and 100 bp ladders were used. (E) PCR confirmation of *galK::gfp* integration into *E. coli* DH5α. (F) Fluorescence microscopy showing enhanced GFP fluorescence observed after *galK::gfp* integration to generate a tester-GFP strain for competition assays.

### 3.5 The EasyOligo strategy for producing donor DNAs

To eliminate precloning steps, a PCR-based strategy using oligonucleotides, akin to the EasyOligo approach for gRNAs, was developed to directly produce donor DNAs. This method is demonstrated by generating a 32 bp deletion in *galK* (Fig. 5A). The deletion is specified within a central 50-nt oligo, amplified by PCR using 60-nt forward and reverse oligos providing chromosomal recombination flanks. The resulting 140 bp dsDNA and the gRNA plasmid are directly transformed into *E. coli* via heat shock for CRISPR/Cas9- mediated deletion (Fig. 5A).

**Fig. 5.**
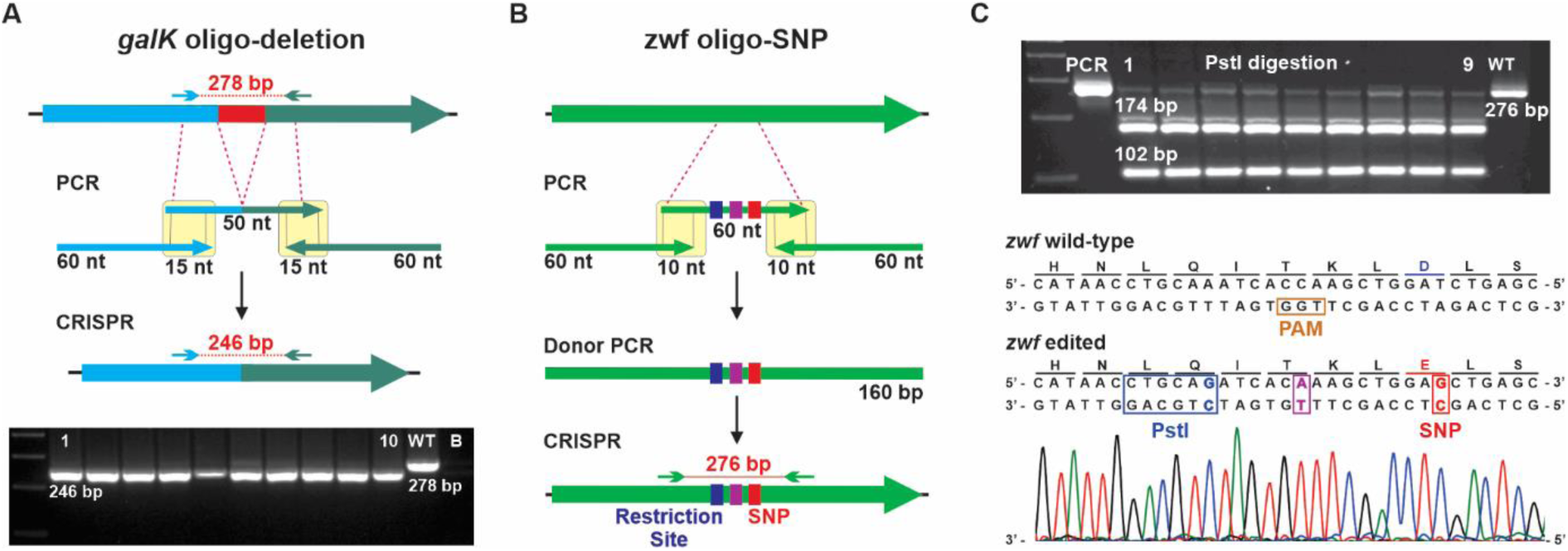
Production of donors using oligonucleotides. (A) Generation of a donor to produce a 32 bp deletion in *galK*. Deletion borders were specified within a 50-nt central oligo, which was used for amplifying a 140 bp PCR product with forward and reverse oligos containing 15-nt priming sites. 2% agarose gel electrophoresis shows a 246 bp fragment for the deletion and a 278 bp band for WT. (B) The editing of SNPs in the *zwf* gene. The *zwf* T/G mutation (red box), the obliteration of the PAM sequence (purple), and the creation of a PstI restriction site (blue) are specified within the 60-nt central oligo, which serves as a template for the PCR amplification of a 160 bp donor, directly transformed into *E. coli* for CRISPR/Cas9 genome editing. (C) Editing confirmed by PstI digestion and Sanger sequencing; cleaved fragments of 174 bp and 102 bp indicate successful editing, while a 276 bp fragment corresponds to WT.

In another example, a 60-mer central oligo was used to reverse-engineer the T\G transversion in the *zwf* gene from the evolved E-Cm population into DH5α (Fig. 5B). A PstI restriction site was incorporated into the oligo to identify correctly edited colonies, and the PAM consensus was altered to prevent Cas9 recognition. These edits did not change the encoded amino acids (Fig. 5C). After transformation, positive colonies were screened using PstI restriction analysis, and DNA modifications were confirmed by Sanger sequencing (Fig. 5C).

Interestingly, attempts to transform oligo-derived PCR products as donors via electroporation failed to produce colonies or resulted in aberrant genomic modifications. Similarly, in vivo cloning of gRNAs was consistently problematic. A comparison of electroporation and heat shock transformations for the deletions of *galK*, *flhCD*, and *hflK* confirmed the drawbacks of electroporation with oligo-derived donors (Fig. S3A-H). These results align with previous reports indicating that in vivo cloning in *E. coli* was more successful with heat shock transformation than with electroporation (Bubeck et al., 1993; Kostylev et al., 2015).

For the *hflK* deletion, a third donor DNA production method was employed involving PCR generation of a repair template with asymmetric arms (Jacobus et al., 2022; Jacobus et al., 2024; Shukal et al., 2022). A minimized 25 bp left flank was utilized. Donor DNA was also directly PCR-amplified from the *E. coli* genome (Table 2).

### 3.6 Reverse engineering of ALE mutations

We utilized our CRISPR EasyGuide approach to reverse engineer a set of alleles from sucrose-evolved populations into *E. coli* DH5α (Table 2). To assess the impact of individual mutations on sucrose growth fitness through competition assays (described below), a testing strain was constructed by inserting the *cscBKA* operon into the DH5α SS9 intergenic region (Bassalo et al., 2016). Despite our efforts, we could not transform the Cas9/λ-Red recombinase plasmid into *E. coli* E2348/69. Therefore, *E. coli* DH5α SS9::*cscBKA* was also used to integrate selected mutations from E2348/69 populations. Genes mutated in different ALE populations, indicative of parallel evolution, are reliable markers of adaptive processes (Menegon et al., 2022). Thus, our reverse engineering campaign prioritized alleles from parallel mutated targets (Figure 2A and B) and mutations potentially affecting growth fitness or sucrose consumption under ALE conditions (Table 2).

Following the CRISPR/Cas9 procedure, we screened the transformants to assess positive editing rates through colony PCR. Overall, the efficiency rates achieved were high, with numerous targeted genomic loci being edited with 100% efficiency (Table 2). This high level of precision underscores the effectiveness of our CRISPR/Cas9 approach. However, it is important to note that some loci exhibited low editing efficiency. This discrepancy in efficiency levels may be attributed to the variable performance of specific gRNAs (Bassalo et al., 2016; Shukal et al., 2022). For instance, the editing of the *lacZ* locus demonstrated lower success rates. These observations highlight the importance of gRNA selection in obtaining optimal editing results. Detailed outcomes of each individual CRISPR/Cas9 experiment are documented in the Supplementary File 1.

### 3.7 Fitness evaluation of evolved populations and engineered strains

We assessed the fitness of the ALE populations and each reverse-engineered strain using a growth competition assay against a GFP-expressing reference strain (Fig. 6A). Flow cytometry was employed to measure the ratio of fluorescent to non-fluorescent cells at the start and end points of propagations, enabling the calculation of a selection coefficient normalized per cell doubling (S/doubling). A positive S/doubling demonstrates that the evolved population or engineered mutant has an enhanced growth capacity (i.e., fitness) compared to the parental strain, while a negative S/doubling indicates that the engineered allele negatively affects growth fitness.

**Fig. 6.**
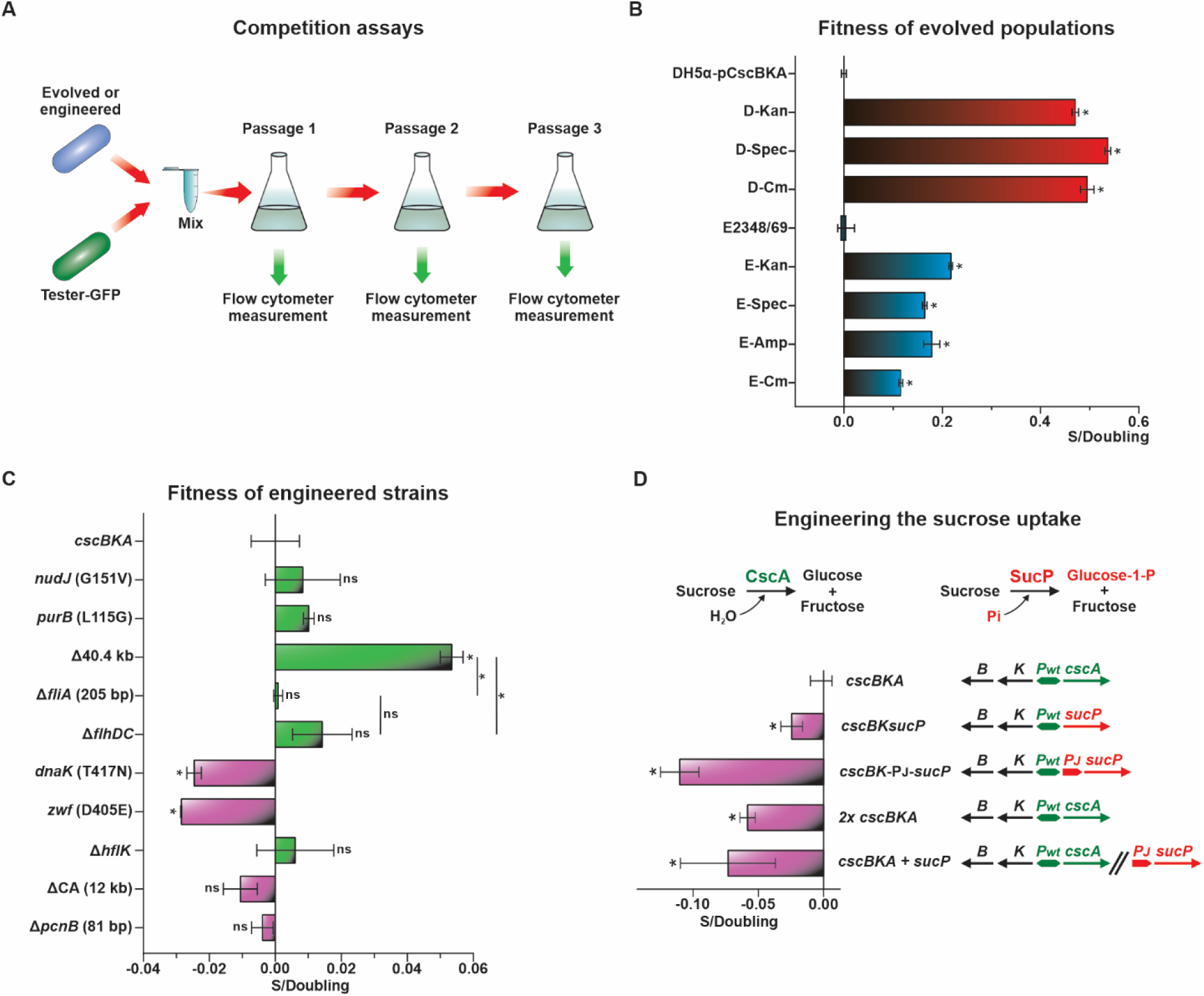
Fitness estimations for evolved populations and engineered strains. (A) Competition experiment setup: strains and tester-GFP were mixed in specific proportions in minimal medium containing 2% sucrose and propagated in triplicate. Flow cytometry measurements of GFP versus non-GFP cells at the initial point and over one to three passages enabled the calculation of selection coefficients per doubling (S/doubling). (B) Fitness for evolved populations, normalized to the progenitors (*E. coli* DH5α-pCscBKA or E2348/69), was calculated following one passage. (C) Fitness for engineered strains was assessed over three passages. (D) Fitness of strains combining the *cscBKA* and *sucP* genes. *Pwt* (green) represents the E2348/69 *cscBKA* operon promoter; *Pj* (red) is the synthetic prJ23119 promoter. S/doubling values were normalized to the value obtained for the competition of the tester-GFP versus *E. coli* DH5α SS9::*cscBKA*. Statistical significance was compared to *E. coli* DH5α SS9::*cscBKA* and among the *fliA*, *flhDC*, and Δ40.4 kb strains. ns: non-significant; (*) significant (*p* < 0.05).

Competitions with the evolved ALE populations revealed increased fitness compared to their evolutionary ancestors (Fig. 6B). The DH5α populations demonstrated a greater fitness improvement than the E2348/69 populations, which aligns with the higher µmax observed in microplate growth assays (Fig. 1B). This fitness boost in DH5α is likely due to the chromosomal integration of the pCscBKA plasmid in the D-Kan and D-Cm populations, along with decreased pCscBKA copy number in D-Spec (Fig. 2C).

Among the reverse-engineered ALE mutations, only the 40,419 bp deletion from the D-Spec population demonstrated a significant positive fitness effect (Fig. 6C). This deleted region encompasses the *fli* operons related to flagellum assembly (Juhas et al., 2014). To explore the fitness increase associated with impaired flagellum assembly, we deleted the *fliA* gene, a late-stage flagellum regulator, and the *flhDC* operon, a primary flagellum regulator (Barembruch and Hengge, 2007; Chilcott and Hughes, 2000). Neither deletion resulted in a fitness increase comparable to that of the 40,419 bp deletion. This gene cluster also includes the *rcsA* ORF (Fig. 2D), a positive regulator of CA biosynthesis (Ebel and Trempy, 1999). A partial deletion of the CA biosynthetic cluster was observed in E2348/69 E-Kan and E-Cm populations, suggesting adaptive parallelism. Testing the CA cluster deletion in DH5α revealed a slightly negative fitness effect (Fig. 6C).

Competing reverse-engineered alleles related to *purB*, *nudJ*, *hflK*, *zwf*, and *dnaK* with the tester-GFP exhibited null or negative fitness effects (Fig. 6C). Similarly, an 81 bp deletion in *pcnB* in the D-Spec population slightly hindered growth. However, the Δ*pcnB* mutation positively impacted fitness in DH5α strains expressing the pCscBKA plasmid (see Fig. 7).

**Fig. 7.**
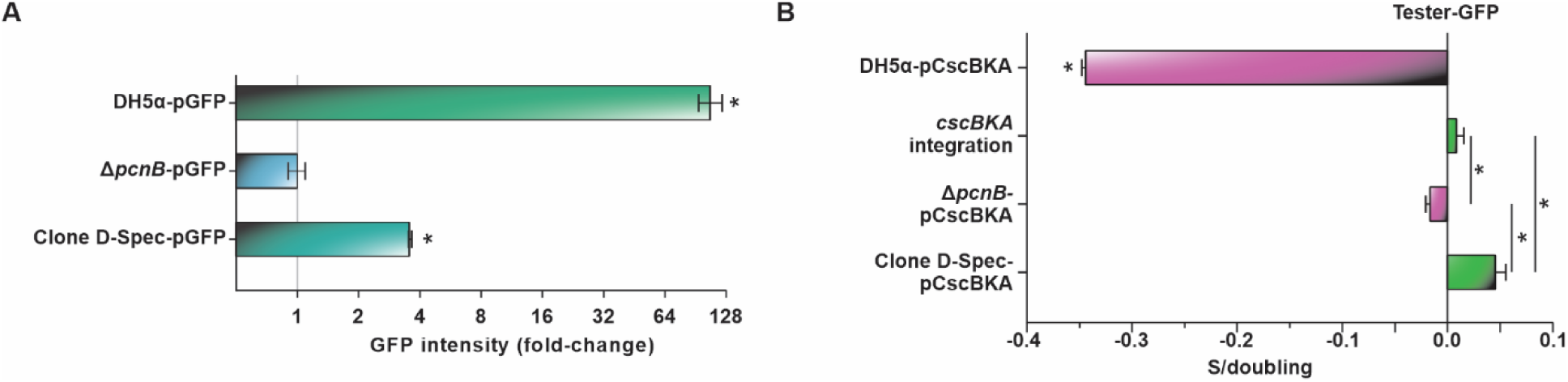
**Impact of mutations on plasmid copy numbers and pCscBKA fitness**. (A) GFP expression as a proxy for estimating relative differences in plasmid copy numbers. The high-copy pGFP plasmid was transformed into *E. coli* backgrounds: DH5α (reference), DH5α Δ*pcnB*, and the evolved clone D-Spec. GFP levels were measured by flow cytometry, with experiments conducted in triplicate. (B) The fitness impact of pCscBKA copy numbers was assessed through competitions with the tester-GFP. The pCscBKA plasmid was transformed into the *E. coli* backgrounds: DH5α, DH5α Δ*pcnB*, and the evolved clone D-Spec. The *cscBKA* integration was into the DH5α SS9 *locus*. (*) Indicates significant differences (*p* < 0.05) between strains.

### 3.8 Engineering operons for sucrose catabolism

We leveraged the CRISPR EasyGuide system to engineer variations in the *cscBKA* operon (Fig. 6D), including replacing *cscA* (invertase) with the *Bifidobacterium adolescentis sucP* ORF, which encodes sucrose phosphorylase (van den Broek et al., 2004). This phosphorolysis pathway spares one ATP compared to sucrose hydrolysis, where glucokinase (Glk) consumes ATP to phosphorylate glucose (Fig. 1A) (Olavarria et al., 2019). We tested four engineered strains to explore the adaptive effects of encoding *sucP* with the *cscBKA* operon. Competition assays in minimal medium with 2% sucrose (Fig. 7D) showed that expressing *sucP* combined with *cscBK* within the operon or in parallel with *cscBKA* reduced fitness compared to *cscBKA* alone. Additionally, expressing two copies of *cscBKA* also decreased growth fitness on sucrose (Fig. 6D). The results show that under our tested conditions a single copy of the *cscBKA* operon is more effective, yielding greater fitness than its duplication or combinations with *sucP*.

### 3.9 Reduced fitness associated with cscBKA expression from plasmids

We observed a significant reduction in pCscBKA copy numbers in the evolved DH5α populations. Potential chromosomal integration of pCscBKA was noted in populations D- Kan and D-Cm, while both pCscBKA and pSpec plasmids in D-Spec population had reduced copy numbers (Fig. 2C). The D-Spec population also contains an 81 bp deletion in *pcnB*, which encodes poly(A) polymerase I (PAP I) and was identified in a screening for mutants that lower ColE1 plasmid copy numbers, hence its name (plasmid copy number B) (Lopilato et al., 1986). To investigate the role of *pcnB* in maintaining ColE1 plasmids in DH5α, we used a plasmid-based GFP reporter (pGFP, a ColE1-type plasmid) for estimating relative differences in plasmid copy numbers. Fluorescence intensity from GFP-expressing plasmids, measured by flow cytometry, correlates with plasmid copy numbers (Joshi et al., 2022). We isolated a colony from the D-Spec population and propagated it to eliminate the pCscBKA and pSpec plasmids. The curated D-Spec clone, the reverse-engineered strain Δ*pcnB*, and DH5α were transformed with pGFP, and fluorescence intensity was measured in triplicate at the stationary growth phase. The D- Spec clone and Δ*pcnB* mutant showed a marked decrease in GFP expression compared to DH5α, consistent with lower pGFP copy numbers (Fig. 7A).

Through competition assays with the tester-GFP, we estimated the fitness associated with plasmid-based (pCscBKA) versus chromosomal expression of *cscBKA* (Fig. 7B). A significant decrease in fitness was observed in DH5α carrying pCscBKA compared to the tester-GFP with genomic integration of *cscBKA*. Our data suggest a fitness drop of approximately 35% per doubling for *cscBKA* expressed from a high-copy plasmid compared to its chromosomal counterpart. This burden was alleviated by the Δ*pcnB* mutation alone or within the D-Spec clone background. Additional mutations, such as the Δ40.4 kb deletion in the D-Spec background, may contribute to the positive fitness observed in the evolved clone. These findings confirm that the integration or reduction in copy number of the *cscBKA* operon was a primary driver of fitness gains in the evolved DH5α populations.

### 3.10 Integration of the phaCAB operon for PHB production

To demonstrate the utility of the EasyGuide approach for the metabolic engineering of *E. coli*, we assembled the *phaCAB* cassette and performed its chromosomal integration into the *E. coli* SS9::*cscBKA* strain. The *phaCAB* operon from *Cupriavidus necator* supports the synthesis of PHB, a polyester-type polymer providing a biobased alternative to oil-derived plastics (Hiroe et al., 2012; Sohn et al., 2020). Producing PHB in *E. coli* strains engineered for sucrose consumption enables the use of cost-effective and renewable feedstocks for industrial applications (Sohn et al., 2020). We utilized yeast in vivo recombination to assemble the *C. necator phaCAB* cassette, transforming a PCR product containing the structural genes and flanking regions into *E. coli* DH5α, targeting the *lacZ* locus for integration. A strong prJ23119 promoter was added to drive *phaCAB* expression (Fig. 8A). PHB production from sucrose in the transformed strain was confirmed through GC-MS analysis, yielding approximately 31% of dry cell mass, close to the 36% produced by *Burkholderia glumae*, a natural PHB producer (Fig. 8B and C).

**Fig. 8.**
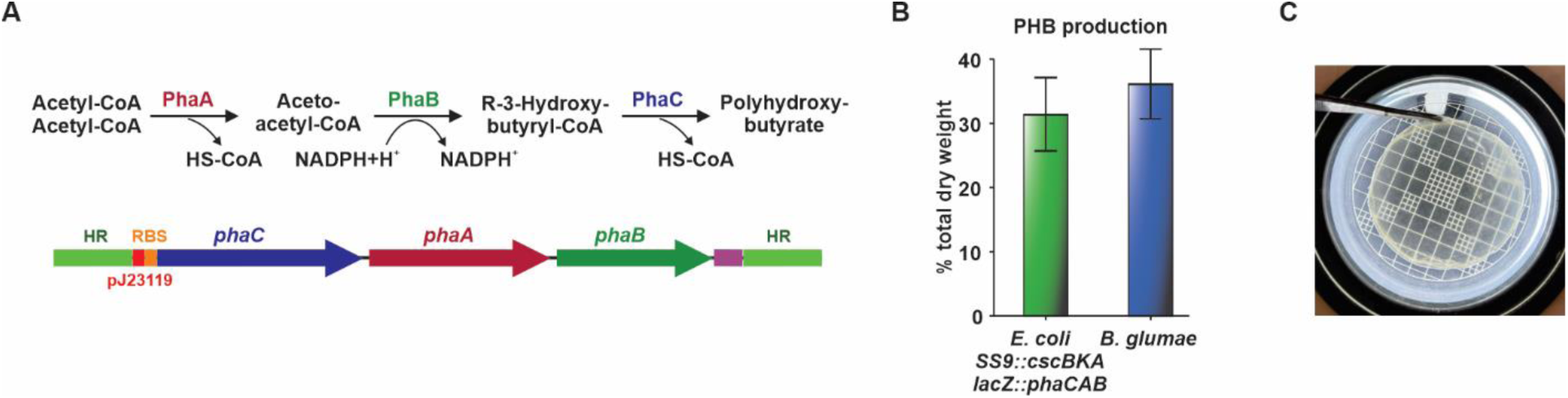
PHB production from sucrose in *E. coli* DH5α with integrated *cscBKA* and *phaCAB* operons. (A) Metabolic pathway for PHB production and the *phaCAB* construct for insertion, assembled in yeast. After cloning, the donor was amplified for insertion into the *lacZ* locus, with expression driven by a strong constitutive promoter (prJ23119). (B) PHB production as a percentage of dry cell mass in engineered DH5α and *B. glumae*. (C) PHB film produced by engineered *E. coli* DH5α.

## 4. Discussion

We conducted an ALE experiment to enhance the growth of *E. coli* on sucrose, resulting in the identification of various mutations, including deletions, insertions, and SNPs across seven populations. To rigorously validate these ALE alleles, we used CRISPR/Cas9-mediated reverse engineering alongside innovative strategies for donor DNA production and the adaptation of the EasyGuide system for efficient gRNA recombination cloning in *E. coli* (Jacobus et al., 2022). Our in vivo methods provide a cost-effective and practical alternative to traditional in vitro gRNA/donor preparation techniques, such as restriction-ligation, CPEC, or Golden Gate assembly (Table 1).

The in vivo recombination pathway in *E. coli* was described in the early 1990s for cloning PCR products (Bubeck et al., 1993; Jones and Howard, 1991; Oliner et al., 1993). Despite numerous applications documented since then (Jacobus and Gross, 2015; Nozaki and Niki, 2019; Watson and Garcia-Nafria, 2019), its potential as a simplified cloning tool has been largely overlooked. In our study, we capitalized on this pathway using three distinct gRNA assembly strategies: (i) Plasmid circularization, which involves PCR amplification of pTargetAmp with primers featuring 5’ overlapping spacers for in vivo plasmid closure. While practical, this method is susceptible to false positives due to template plasmid carryover during transformation (Jacobus and Gross, 2015). (ii) Bimolecular complementation effectively reduces false positives. Cotransforming two PCR products from pSpec5’ and pSpec3’ enables the formation of spectinomycin-resistant colonies if precise recombination assembles both the gRNA and the complete *aadA* gene. This minimizes false positives, as shown in the in vivo assembly of split yeast episomes (Jacobus et al., 2022; Kuijpers et al., 2013). (iii) The EasyOligo approach simplifies the process by using PCR amplification of a single 50-mer oligonucleotide containing the gRNA spacer; the resulting 140 bp dsDNA product is easily cotransformed with pre-prepared pOliSpec5’ and pOliSpec3’ fragments to generate SpecR colonies.

The production of editing templates for CRISPR applications in *E. coli* has traditionally been limited by the need to preassemble large HA (100–600 bp) for high editing efficiencies (Bassalo et al., 2016; Jiang et al., 2015). To overcome this challenge, we repurposed our pEasyG3 plasmids (Jacobus et al., 2022) to enable in vivo yeast fusion of PCR fragments flanking deletion or editing sites and to assemble insertion cassettes. The time invested in yeast transformation, colony screening, and DNA preparation is compensated by exceptionally high assembly rates, often reaching up to 100%.

As an alternative to preassembling donors, synthetic oligonucleotides (40–120-mer) serve as effective donor DNAs, demonstrating efficient recombineering rates even for large genomic deletions, with double-stranded oligos outperforming single-stranded ones (Reisch and Prather, 2015; Zerbini et al., 2017). We also used these oligos as templates for PCR production of dsDNA donors with up to 70 bp HA, embedding the editing sequence or deletion borders within the Mid-oligo. This practical approach is streamlined by a thermal-shock transformation protocol, which eliminates the need for PCR product desalting/purification, a required step for electroporation in typical *E. coli* CRISPR applications. This simplified protocol allows direct transfer of DNA from PCR tubes into chemically competent cells.

By employing the EasyGuide approach with yeast-assembled and oligo-derived donors, along with repair templates that were PCR-amplified from ALE populations, we consistently achieved high editing rates, often up to 100% positive outcomes. However, as noted in various studies (Bassalo et al., 2016; Shukal et al., 2022), editing efficiencies varied among different gRNAs, with targets such as *lacZ*, *pcnB*, and *dnaK* showing positive rates as low as 10%. Overall, EasyGuide proved to be a robust method for reverse engineering multiple mutations identified in ALE populations propagated on 2% sucrose. This approach allowed us to assess numerous alleles through competition assays, revealing positive fitness effects for SS9::*cscBKA*, Δ*pcnB*, and the ∼40.4 kb deletion that includes the flagellar assembly cluster. A significant fitness decrease of approximately 35% per cell doubling was observed for plasmid-based expression of *cscBKA* compared to SS9::*cscBKA*, indicating that the evolution of D-Kan and D-Cm populations was primarily driven by the chromosomal integration of pCscBKA. Similarly, the fitness improvement in the D-Spec population likely stemmed from reduced pCscBKA copy numbers due to Δ*pcnB*. In *E. coli* K-12, decreased growth rates linked to *cscBKA* plasmids were also alleviated by the operon integration into the genome (Bruschi et al., 2012; Carruthers et al., 2020).

A reduced dosage of CscBKA may alleviate its metabolic burden, while the deletion of a 40,419 bp segment in D-Spec, which includes flagellar assembly regions 3a and 3b (Juhas et al., 2014), likely impacts the cell’s energy dynamics. Flagellar assembly in *E. coli* requires the expression of over 60 genes (Barembruch and Hengge, 2007; Chilcott and Hughes, 2000), consuming approximately 5.0% of the cell’s energy, with an additional 5.2% dedicated to flagellar rotor operation (Schavemaker and Lynch, 2022). Thus, the observed ∼5% fitness gain per doubling from deleting flagellar clusters is expected; however, this improvement exceeds that achieved by individually deleting the key flagella regulators *flhDC* and *fliA* (Barembruch and Hengge, 2007; Chilcott and Hughes, 2000). Other genes within the 40,419 bp segment may also contribute to adaptive effects, such as *rcsA*, a transcription factor involved in CA biosynthesis (Ebel and Trempy, 1999). However, partial deletion of the 12.0 kb CA cluster and additional mutations in *E. coli* DH5α and E2348/69 have shown negative or neutral impacts on fitness in DH5α under 2% sucrose conditions. The lack of notable adaptive effects from reverse-engineered ALE alleles may stem from their specific genetic backgrounds or epistatic interactions (Barrick and Lenski, 2013; Menegon et al., 2022). For instance, recreating the loss-of-function mutation in *hflK* from E2348/69 in DH5α yielded only a negligible fitness gain. Mutations in *hflK* and *hflB*, as observed in E2348/69, have been associated with promoting λ prophage dormancy (Bandyopadhyay et al., 2011), likely helping to mitigate genome instability in the prophage-carrying strain E2348/69. Since the DH5α genome lacks λ prophages, the *hflK* deletion may not confer any adaptive benefits for this strain.

ALE with DH5α populations identified mutations in *rpoB*, *rpoC*, and *rpoZ*, which have been linked to enhanced growth rates in minimal media utilizing glucose or sucrose (Conrad et al., 2010; Mohamed et al., 2019). These findings, along with other mutations, suggest that our ALE strategy favored systemic changes that improved overall growth fitness rather than specifically optimizing sucrose utilization pathways. Nonetheless, our study provides a basis for advancing sucrose-utilizing platforms. The competitive fitness observed from sucrose hydrolysis (via CscA) compared to phosphorolysis (via SucP) highlights the need to evaluate various sucrose uptake mechanisms and explore optimized combinations. Furthermore, robust stress tolerance is essential for industrial applications and can be enhanced through ALE (Dragosits and Mattanovich, 2013; Menegon et al., 2022). An ideal *E. coli* sucrose-utilizing platform would integrate growth fitness, effective sugar uptake mechanisms, and increased molasses tolerance, facilitating the efficient production of biobased chemicals and fuels (Kim et al., 2024; Mohamed et al., 2019; Olavarria et al., 2019). In this context, we achieved promising results by enabling PHB production from sucrose in *E. coli* using EasyGuide CRISPR, with yields comparable to that of a naturally producing strain. Ultimately, our study demonstrates that combining CRISPR-mediated genetic engineering with ALE is crucial for unlocking the full catabolic potential of microbial chassis.

## Authors contributions

**Joneclei A Barreto:** Writing – original draft, Visualization, Investigation, Formal analysis, Data curation, Methodology. **Matheus V M L e Silva:** Methodology, Investigation. **Danieli C Marin:** Methodology, Investigation. **Michel Brienzo:** Methodology, Investigation, Writing – review & editing. **Ana Paula Jacobus:** Methodology, Conceptualization, Writing – review & editing, Funding acquisition. **Jonas Contiero:** Methodology, Investigation, Conceptualization, Writing – review & editing, Funding acquisition. **Jeferson Gross:** Conceptualization, Visualization, Investigation, Formal analysis, Data curation, Methodology, Writing – original draft, Funding acquisition.

## Data availability

Illumina sequence reads are available at the NCBI under the BioProject number PRJNA1200122.

## Supporting information

Supplementary File 1_ molecular genetic procedures

## Acknowledgements

E2348/69 strain was kindly donated by Dr. Fernanda Batista de Andrade (Instituto Butantan, São Paulo, Brazil). We also thank Prof. José Gregório Cabrera Gomez (University of São Paulo, Brazil) for donating the pBBR1MCS-2 plasmid containing the *Cupriavidus necator phaCAB* operon. This work was supported by grants from the São Paulo Research Foundation numbers 17/50249-6, 20/02246-0, 17/22401-8, 17/24453-5, 23/04162-7, and the Brazilian National Council for Scientific and Technological Development (CNPq) grant 404603/2023-8.

## Appendix A. Supplementary data

Supplementary data to this article can be found online at

