## Supplementary Figs. 1-10 for "Engineering Adaptive Alleles for *Escherichia coli* Growth on Sucrose Using the EasyGuide CRISPR System"

### Engineering Adaptive Alleles for *Escherichia coli* Growth on Sucrose Using the EasyGuide CRISPR System

**Joneclei Alves Barreto[a,b], Matheus Victor Maso Lacôrte e Silva[a,c], Danieli Canaver Marin[a,b], Michel Brienzo[a,b], Ana Paula Jacobus[a,b], Jonas Contiero[a,b,c], Jeferson Gross[a,b]***

[a]*São Paulo State University (Unesp), Institute for Research in Bioenergy, Rio Claro, 13500-230, SP, Brazil*

[b]*PhD Program in Bioenegy, São Paulo State University (Unesp), Rio Claro, 13500-230, Brazil*

[c]*São Paulo State University (Unesp), Institute of Biosciences, Rio Claro, 13506-900, SP, Brazil*

**pTargetAmp**

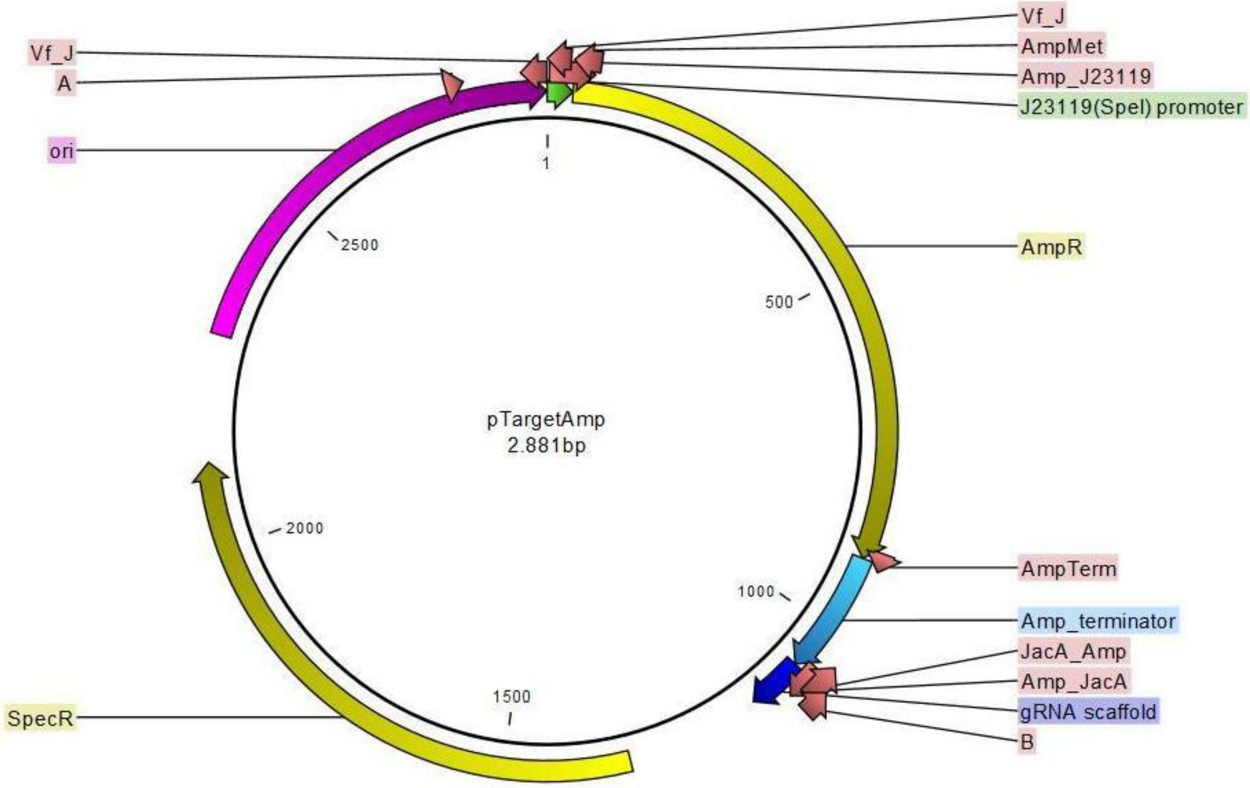


**Supplementary Fig. 1.** pTargetAmp plasmid for the gRNA cloning strategy of plasmid circularization. The prJ23119 promoter is shown in green, and the gRNA scaffold in dark blue. For gRNA plasmid cloning, ori, SpecR, gRNA scaffold, and prJ23119 are amplified by PCR using 5'-end homologous primers.

**pSpec5’**

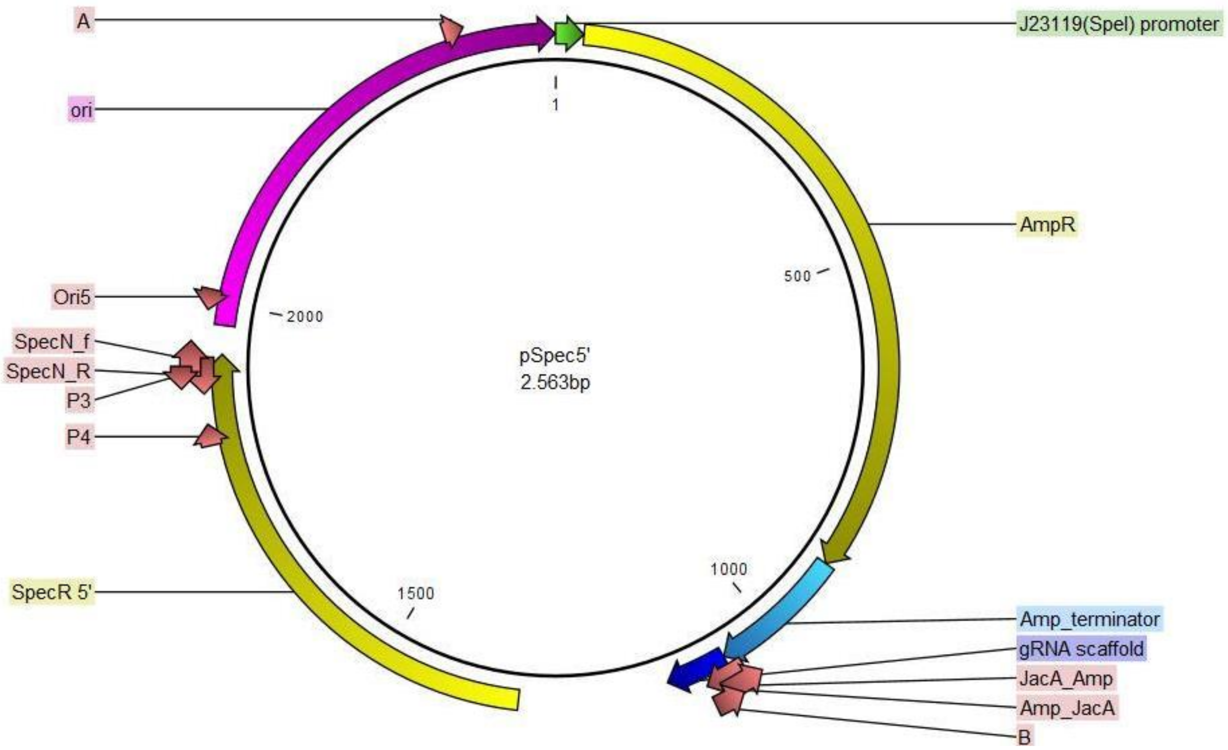


**Supplementary Fig. 2.** Map of the pSpec5’ plasmid. First plasmid used in the bimolecular complementation strategy for gRNA cloning. For gRNA plasmid cloning, the SpecR 5’-end and gRNA scaffold are amplified by PCR with 5’-end primers gRNA and P3.

**pSpec3’**

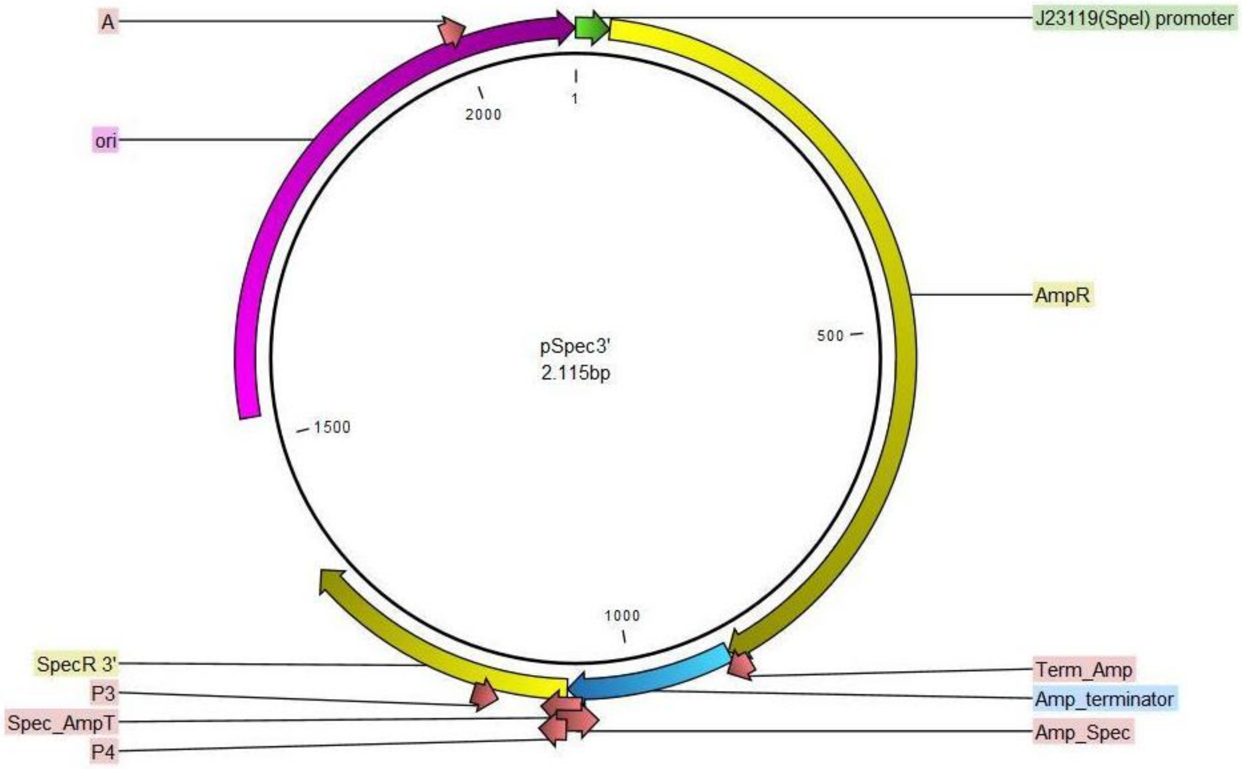


**Supplementary Fig. 3.** pSpec3’ map. Second plasmid used for the bimolecular complementation gRNA cloning strategy. For gRNA plasmid cloning, the SpecR 3’-end, ori, and prJ23119 are amplified by PCR with P4 and 5’-end gRNA primers.

**pOliSpec5’**

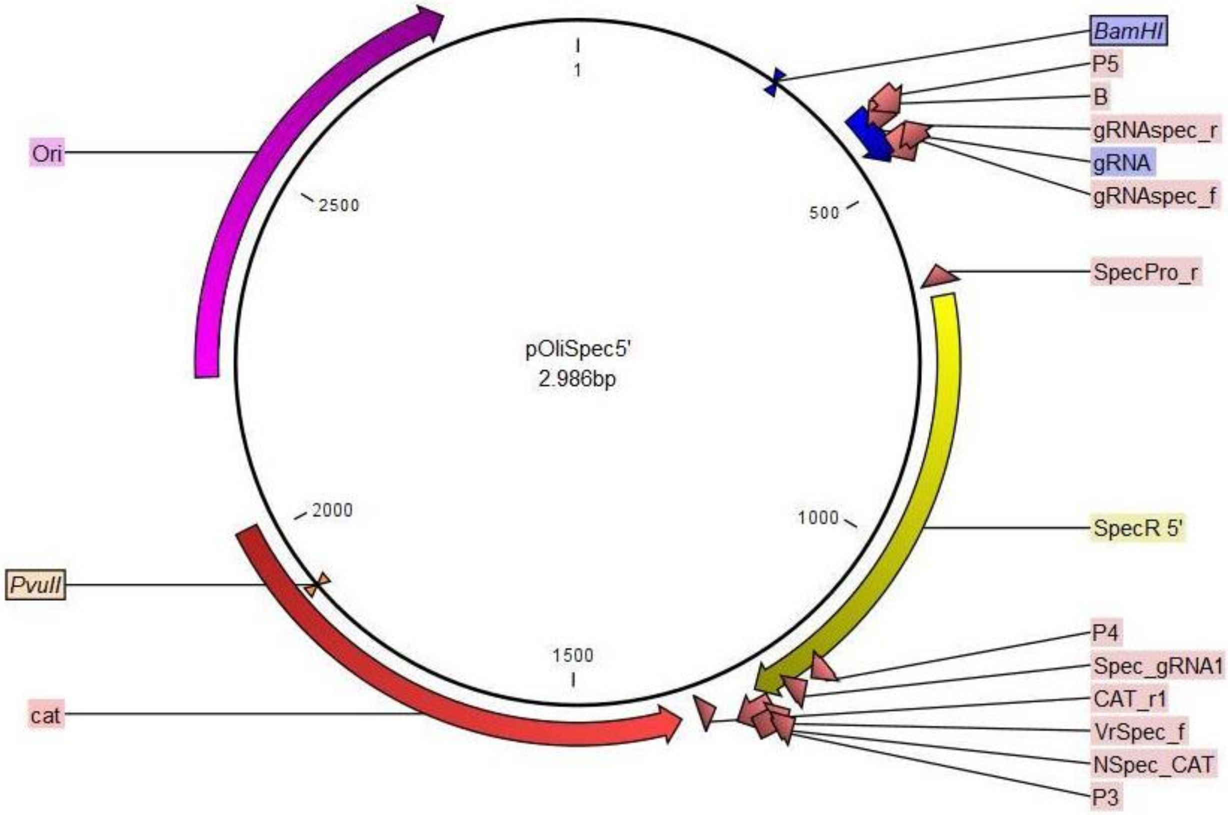


**Supplementary Fig. 4.** pOliSpec5’ map. First plasmid for oligo gRNA cloning. For gRNA plasmid cloning, the SpecR 5’-end and gRNA scaffold are amplified by PCR using P5 and P3 primers, after restriction with BamHI and PvuII enzymes.

**pOliSpec3’**

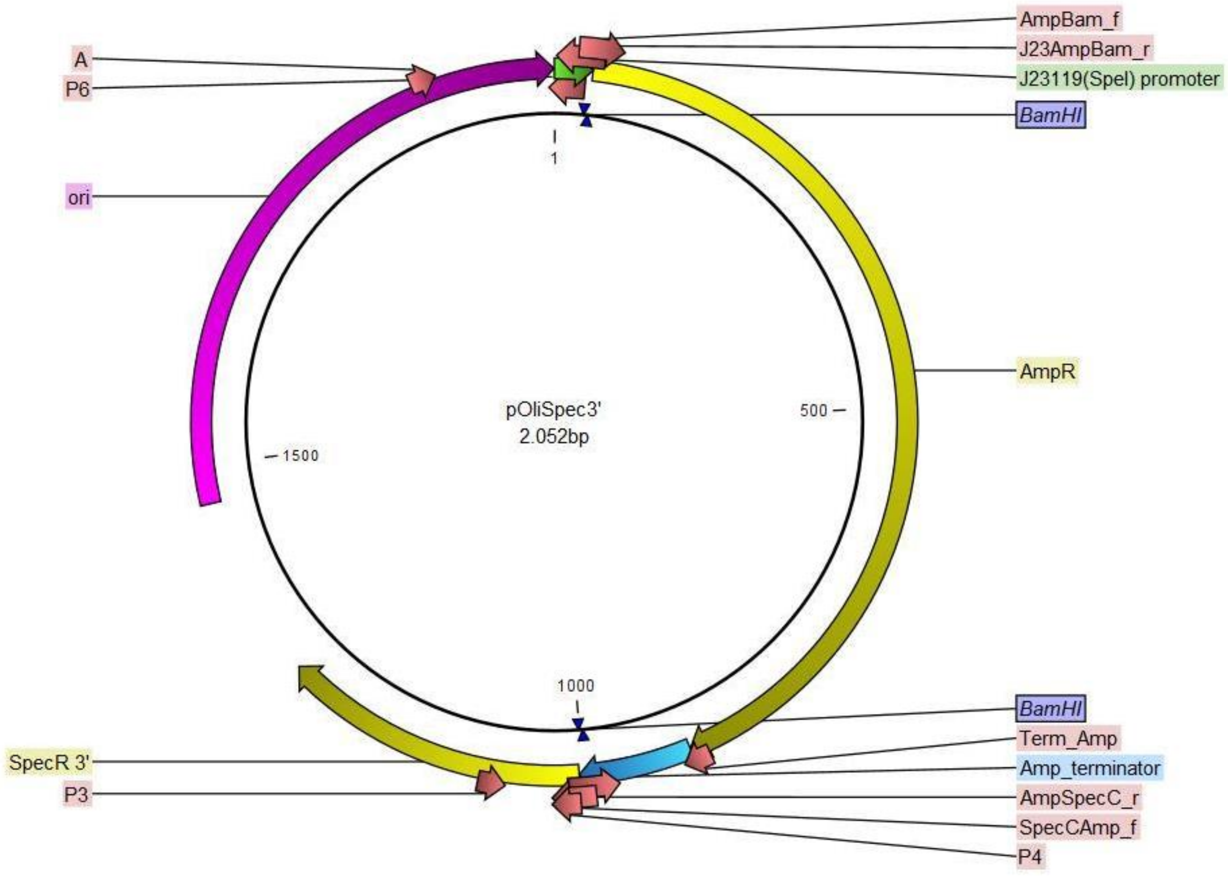


**Supplementary Fig. 5.** pOliSpec3’ map. Second plasmid for oligo gRNA cloning. For gRNA plasmid cloning, the SpecR 3’-end, ori, and prJ23119 are amplified by PCR using P4 and P6 primers, after restriction with BamHI enzyme.

**pCscBKA**

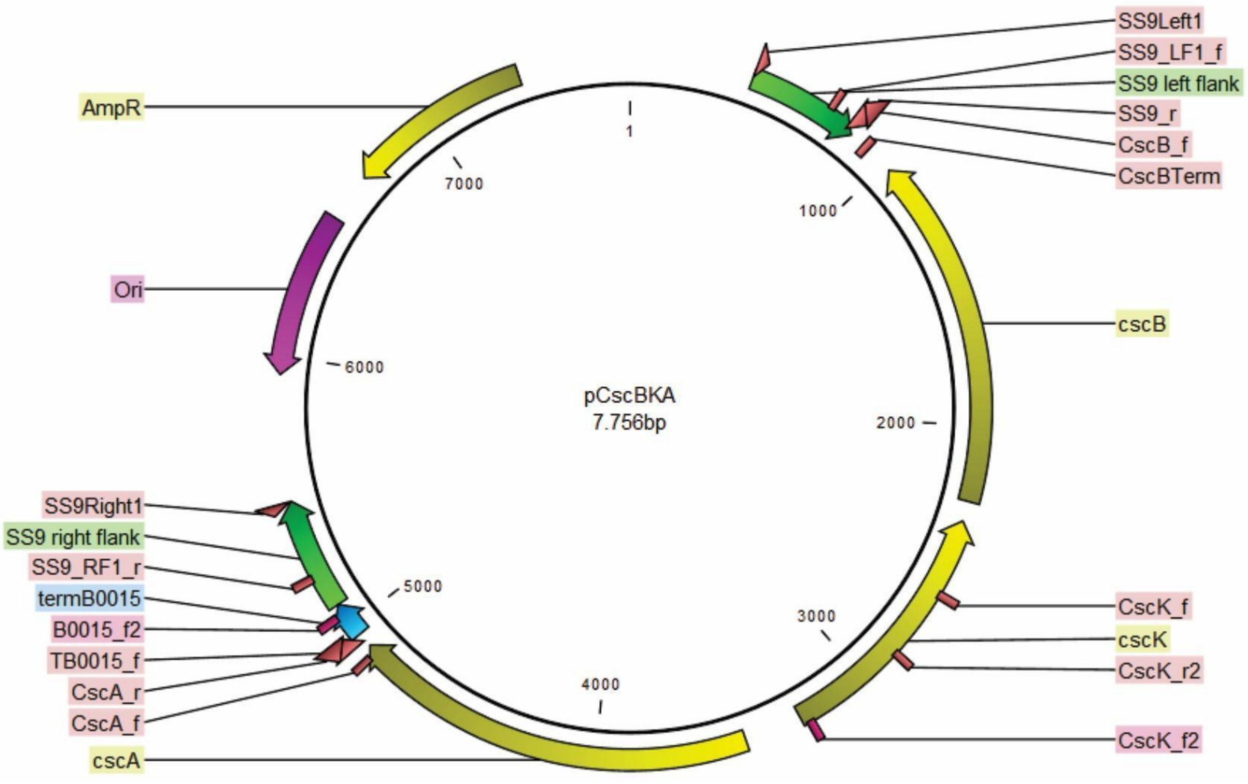


**Supplementary Fig. 6.** pCscBKA map. Plasmid for *cscBKA* operon expression. The SS9 flanking regions are shown in green. For use as a donor, flanking regions and the *cscBKA* operon are amplified with primers SS9Left1/SS9Right1.

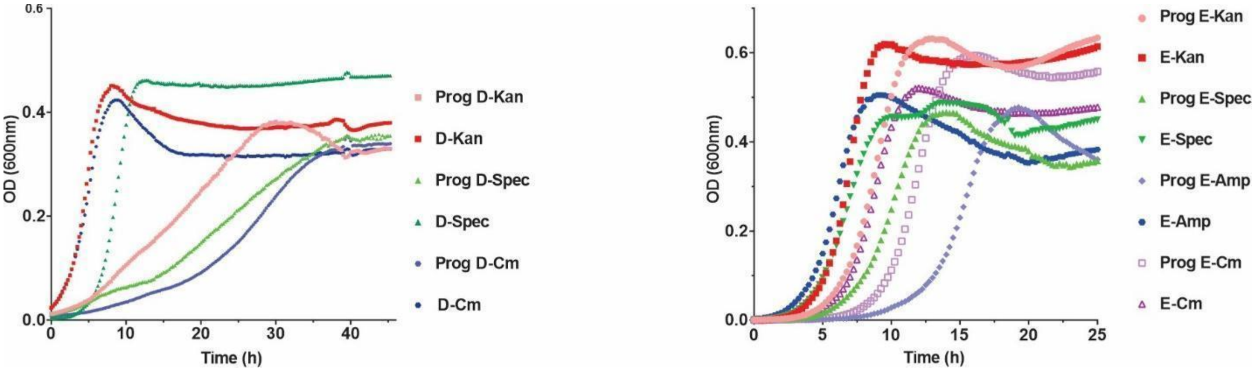


**Supplementary Fig. 7.** Growth curves of progenitor (Prog) and evolved populations of DH5α and E2348/69 on sucrose as the sole carbon source, conducted in a microplate assay.

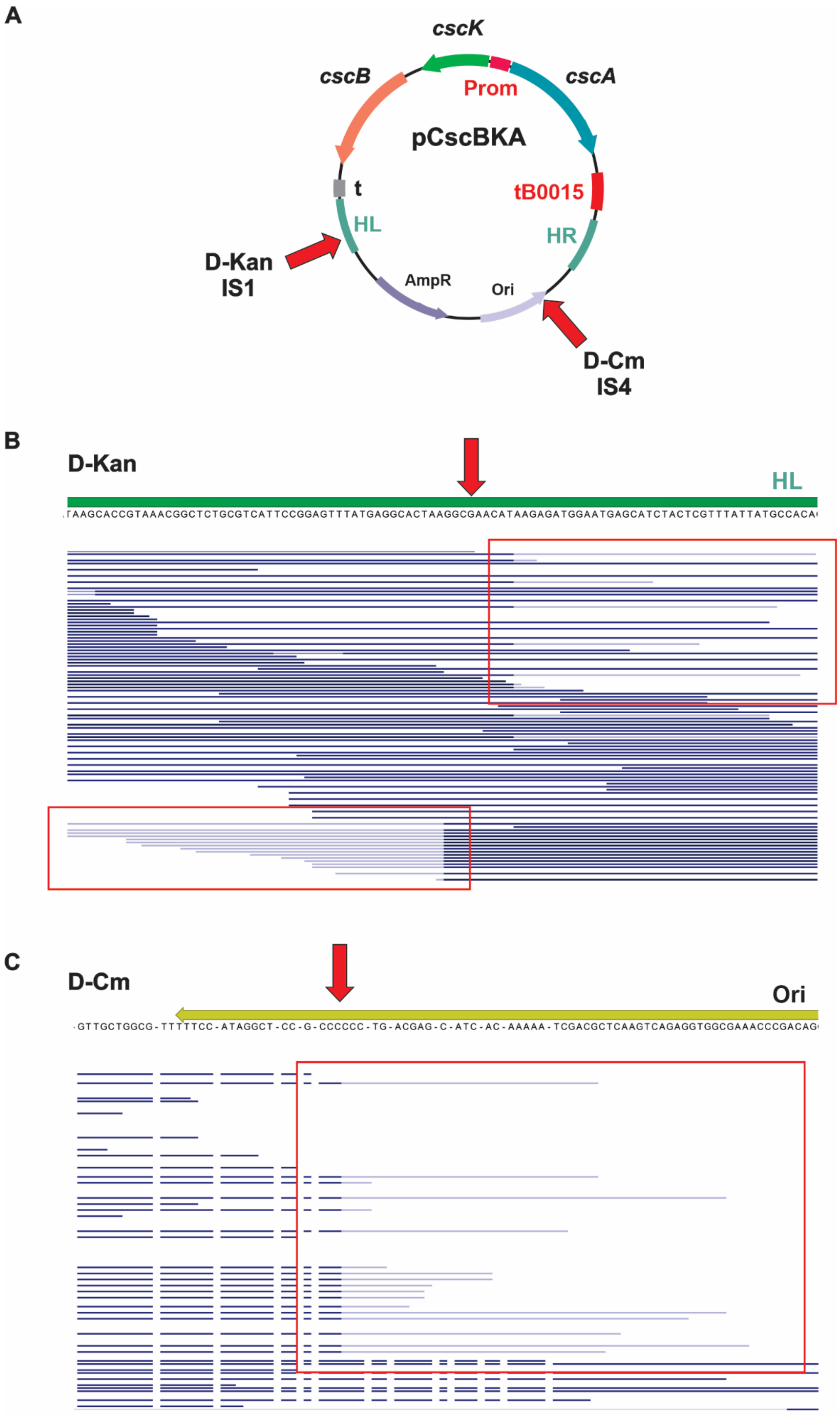
A
cscK
cscB
cscA
Prom
pCscBKA
t
tB0015
HL
HR
D-Kan
IS1
AmpR
Ori
D-Cm
IS4
B
D-Kan
HL
TAAGCACCGTAAACGGCTCTGCGTCATTCCGGAGTTTATGAGGCACTAAGGCGAACATAAGAGATGGAATGAGCATCTACTCGTTTATTATGCCACA
C
D-Cm
Ori
GTTGCTGGCG-TTTTTCC-ATAGGCT-CC-G-CCCCCC-TG-ACGAG-C-ATC-AC-AAAAA-TCGACGCTCAAGTCAGAGGTGGCGAAACCCGACAG

**Supplementary Fig. 8.** Illumina read mappings showing connections between the pCscBKA and *E. coli* Insertion Sequencies (IS). (A) Structure of the pCscBKA plasmid with red arrows showing where Illumina reads map partially presenting unaligned red ends with sequence identities with IS1 (in the D-Kan population) and IS4 (in the D-Cm population). HR and HL are the Homology arms at the Right and Left flanks, respectively, of the *SS9* locus. (B) Read mapping for the D-Kan population showing the region with connection to the IS1. Unaligned read ends are highlighted by the red square. (C) Read mapping for the D-Cm population showing the region with connection to the IS4.

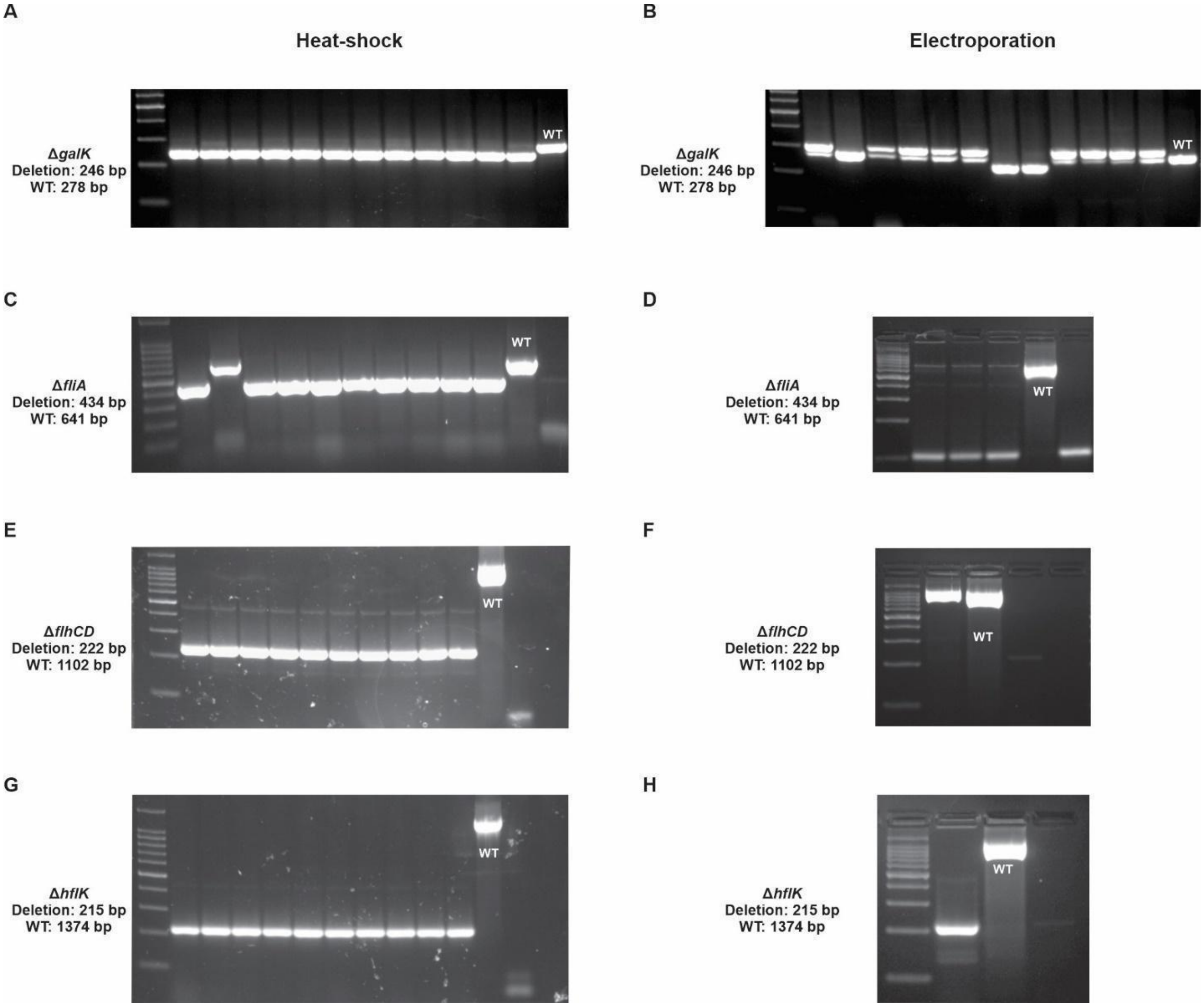


**Supplementary Fig. 9.** Comparison of transformation methods for deletions using heat shock (A, C, E, G) and electroporation (B, D, F, H) for genes *galK*, *fliA*, *flhCD*, and *hflK*. An oligo donor was used for *galK* and *flhCD* deletions, while a donor with asymmetric homology arms was used for *fliA* and *hflK*. In the electroporation transformations for *fliA*, *flhCD*, and *hflK*, all resulting colonies were tested.

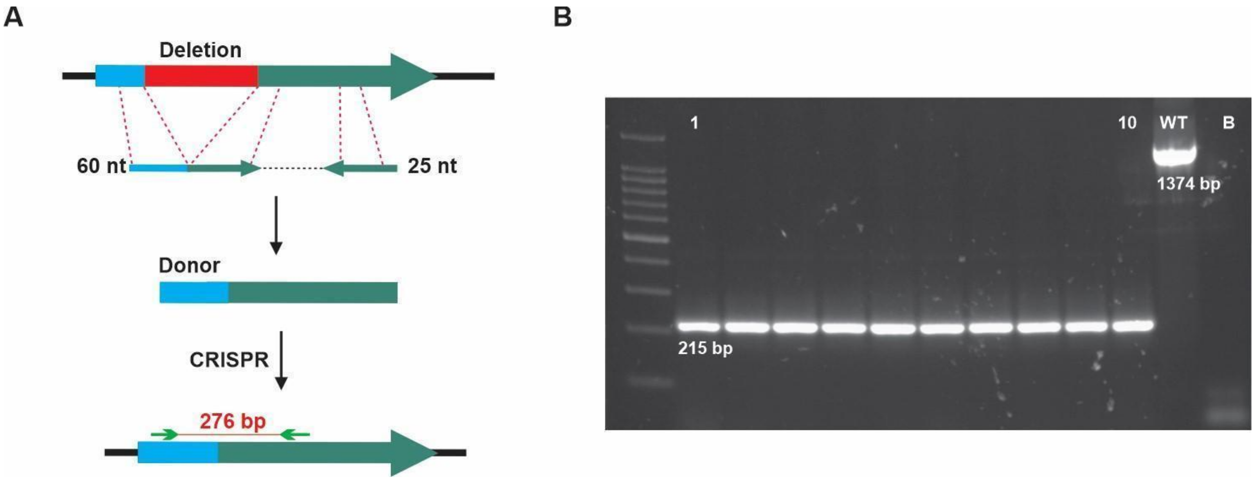


**Supplementary Fig. 10.** Deletion using a donor with asymmetric arms. (A) Structure of gene deletion with an asymmetric donor. To amplify the donor by PCR, one primer specifies the deletion, while a shorter-range primer is used. The homology arms, though unequal, enable recombination to delete the region specified by the longer primer. (B) Gel electrophoresis (2% agarose) showing PCR results of *hflK* deletion in 10 colonies. Positive deletions show a 215 bp band, while wild type (WT) remains intact.
