## Supplementary File 1_ molecular genetic procedures for "Engineering Adaptive Alleles for *Escherichia coli* Growth on Sucrose Using the EasyGuide CRISPR System"

**SUPPLEMENTARY FILE 1: Molecular genetics procedures**

### Engineering Adaptive Alleles for *Escherichia coli* Growth on Sucrose Using the EasyGuide CRISPR System

**Joneclei Alves Barreto[a,b], Matheus Victor Maso Lacôrte e Silva[a,c], Danieli Canaver Marin[a,b], Michel Brienzo[a,b], Ana Paula Jacobus[a,b], Jonas Contiero[a,b,c], Jeferson Gross[a,b]***

[a]*São Paulo State University (Unesp), Institute for Research in Bioenergy, Rio Claro, 13500-230, SP, Brazil*

[b]*PhD Program in Bioenegy, São Paulo State University (Unesp), Rio Claro, 13500-230, Brazil*

[c]*São Paulo State University (Unesp), Institute of Biosciences, Rio Claro, 13506-900, SP, Brazil*

#### gRNA Plasmids cloning

##### Cloning pTargetAmp

The ampicillin resistance marker and terminator were amplified via PCR using primers Amp_J23119/Amp_JacA from plasmid pUC57_sucP, generating a 1,082 bp fragment. The J23119 promoter, origin of replication, spectinomycin resistance marker, and gRNA scaffold were amplified by PCR from plasmid pTargetF using primers JacA_Amp/Vf_J, producing a 1,844 bp fragment. After DpnI treatment, 200 ng of each fragment were mixed and used to transform *E. coli* DH5α cells, using in vivo cloning approach. Colonies were screened with colony PCR using primers A/AmpMet (200 bp).

For use in gRNA plasmid cloning, amplification is performed with primers P2_xxxx/P1_xxxx (1870 bp). where the gRNA sequence is specified by P2_xxxx, and the complementary sequence by P1_xxxx. After treatment with DpnI, the fragment produced was transformed into competent *E. coli* DH5α cells. Colony selection on spectinomycin was followed by confirmation via colony PCR with primers A/B (221 bp).

##### pSpec5' and pSpec3’ cloning and use for plasmid gRNA

For pSpec5' cloning, the ampicillin resistance marker and the origin of replication were amplified via PCR with primers SpecN_f/B (1750 bp) from pTargetAmp. The initial part of the spectinomycin resistance marker and gRNA scaffold were amplified with primers SpecN_R/JacA_Amp (884 bp) from pTargetAmp. After DpnI treatment, the PCR products were mixed and transformed into competent *E. coli* DH5α cells. Positive clones, selected on ampicillin plates, were confirmed by colony PCR with Ori5/P4 primers (182 bp).

For pSpec3' cloning, the ampicillin resistance mark was amplified via PCR with primers Amp_J23119/Amp_Spec (1076 bp) from pTargetAmp, while the terminal part of the spectinomycin resistance marker, replication origin, and J23119 promoter were amplified using Spec_AmpT/AmpMet primers (1,132 bp) from pTargetAmp. After DpnI treatment, with DpnI, the PCR products were mixed and transformed into competent *E. coli* DH5α cells. Positive clones, selected on ampicillin plates, were confirmed via colony PCR with Term_Amp/Spec_gRNA1 primers (247 bp).

For gRNA plasmid cloning, PCR amplification is performed from pSpec5' with primers P3/P2 _xxxx (888 bp); PCR amplification is performed from plasmid pSpec3' with primers P1_xxxx/P4 (1114 bp). The gRNA is specified in primer P2_xxxx and the complementary sequence in primer P1_xxxx. These fragments are mixed and transformed into competent *E. coli* DH5α cells. After colony selection in spectinomycin, cloning confirmation is through colony PCR with the primers A/B (221 bp).

**pOliSpec5' and pOliSpec3' cloning and use for plasmid gRNA**

For pOliSpec5' cloning, the gRNA scaffold and the initial part of the spectinomycin resistance marker were amplified via PCR using the gRNAspec_f/NSpec_CAT_r primers r(832 bp) from pSpec5'. The chloramphenicol resistance marker and origin of replication were amplified from pEasyG3_mic by PCR, followed by DpnI treatment. These PCRs products were mixed and transformed into competent *E. coli* DH5α cells. After colony selection on chloramphenicol plates, positive clones were selected by colony PCR with P4/CAT_r1 (169 bp) P5/SpecPro_r (246 bp) primers.

For pOliSpec3' cloning, the terminal part of spectinomycin, replication origin, and J23119 promoter were amplified by PCR using SpecCAmp_f/J23AmpBam_r (1109 bp) primers from pTargetAmp. The ampicillin resistance and gRNA scaffold were amplified by PCR from pTargetAmp with AmpBam_f/AmpSpecC_r (922 bp) primers. The PCR products were DpnI-treated, mixed, and transformed into competent *E. coli* DH5α cells. After colony selection on ampicillin plates, positive clones were selected by colony PCR with P3/Term_Amp (213 bp) primers.

Prior to amplification for use in gRNA cloning, plasmid pOliSpec5' is digested with PstI and PvuII and pOliSpec3' is digested with BamHI. PCR amplification from digested pOliSpec5' used P5/P3 primers (858 bp) and from pOliSpec3', P4/P6 primers (1,084 bp). These fragments are purified and stored. The gRNA-oligo is amplified by PCR from the central oligo with the primers pE/gE. These three fragments are mixed and transformed into competent *E. coli* DH5α cells. After colony selection in spectinomycin, cloning confirmation is through colony PCR with the A/B (221 bp) primers.

**Cloning plasmids for tagging ALE populations.**

For pKan, two fragments were amplified by PCR from pUC57_phaB using the primers combinations Vf_kan/AmpT_f (641 bp) and pGEX_kan/Or2_r (1,280 bp).

For pSpec, two fragments were amplified by PCR from pTargetAmp, with the primer combinations Vf_spec/AmpT_f (641 bp) and spec_pro/Or2_r (1,253 bp).

For pAmp, a single fragment was amplified from pUC19 using Vf_min/Vr_min primers (1,751 bp).

The PCR products were DpnI-treated and purified. For pKan and pSpec, 200 ng of each fragment were mixed and this mixture was used to transform competent *E. coli* DH5α cells, which were plated on LB plates with the respective antibiotic. For pAmp, 200 ng of the single fragment were used to transform *E. coli* DH5α cells, plated on LB with ampicillin. Colonies were verified by PCR: pKan: prokan_f/Ori_Term_PhaB_R, 164 bp; pSpec: Ori_Term_PhaB_R/prospec_R, 210 bp; pAmp: AmpMet/Ori_Term_PhaB_R, 205 bp.

**Cloning pUC19_B0015 and pGFP**

The Gibson (NEB) method was used to clone pUC19_B0015. The B0015 terminator was amplified by PCR from plasmid pUC57_sucP using B0015for/B0015rev (173 bp) primers, while the vector fragment was amplified from pUC19 using Vf_B0015/pUC19_B0015 primers (2,649 bp). Both products were treated with DpnI. The Gibson followed the manufacturer's instructions, and the product was used to transform competent *E. coli* DH5α cells by heat shock, followed by plating on LB with ampicillin. Colonies were screened by PCR with B0015for/AmpPro (221 bp) primers.

For GFP expression plasmid cloning, the mEGFP sequence was amplified from mEGFP_pBAD plasmid using GFP19for/GFP_B15rev (752 bp) primers. The backbone, containing the lac promoter and B0015 terminator, was amplified from the pUC19_B0015 B15-GFPfor/19GFPrev primers. These fragments were transformed into DH5α cells for cloning and selected for ampicillin. Positive cloning was evaluated using colony PCR using GFP_f/B0015rev (271 bp) and pUC19-2for/GFP_r2 (354 bp) primers. The colonies were cultured, and GFP expression was analyzed by flow cytometry.

#### Donor assembly and CRISPR/Cas9-mediated genomic modifications

##### pCscBKA cloning and SS9::cscBKA insertion

The *cscBKA* operon was amplified by PCR from *E. coli* E2348/69 genomic DNA of using the CscB_f/CscA_r (4,178 bp) primers. The cloning vector, pUC57_SucP, was amplified with SS9_r/TB0015_f (3,695 bp) primers, followed by DpnI treatment. The PCR products were purified, and 200 ng of each were mixed to transform competent *E. coli* DH5α cells, which were plated on LB with ampicillin. Colonies were screened by PCR using SS9_LF1_f/cscB_Term (202 bp) and CscA_f/SS9_RF1_r (362 bp) primers.

From this cloned plasmid, the donor was amplified by PCR for genome insertion of the *cscBKA* operon using the primers SS9Left1/SS9Right1 (5046 bp) purified and 400 ng was used for the CRISPR/Cas9 technique. The insertion of the operon was confirmed by colony PCR with FE1/CscBTerm (582 bp) and B0015_f2/FD1 (524 bp) primers were used. The gel for confirmation of this insertion is represented in Fig. S1.

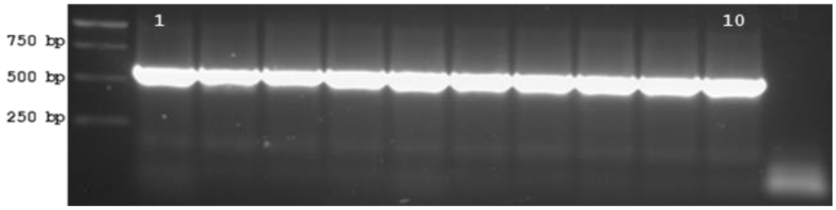


**Fig. S1.** Colony PCR for confirmation of insertion of *cscBKA* operon in the SS9 site. Primers: FE1/CscBTerm (582 bp). The numbers (1-10) indicate the colonies tested.

##### *ΔpcnB* (81 bp)

The donor was amplified by PCR with primers Donor_pcnB_F/donor_pcnB_R (739 bp) from genomic DNA of D-Spec population. and used in CRISPR/Cas9 with the gRNA plasmid in *E. coli* DH5α cells containing the *cscBKA* operon. Deletion was confirmed by colony PCR with pcnB_F/pcnB_R (228 bp for deletion; wild type: 309 bp) primers. Confirmation of this deletion is shown in Fig. S12.

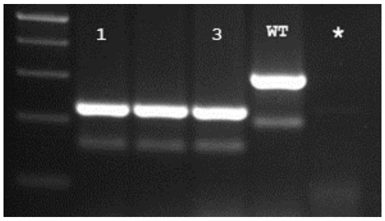


**Fig. S2.** PCR from genomic DNA for confirmation of deletion of 81 bp of *pcnB* gene. Primers: pcnB_F/pcnB_R (228 bp for deletion; wild type: 309 bp).The numbers (1-3) indicate the colonies tested. (*) represents blank; WT: wild type. 100 bp ladder.

## Δ40.4 kb

Primers for PCR amplification of the donor for a 40.4 kb deletion (Flagel_left_F/Flagel_right_R) were designed 1,000 bp upstream and downstream of the target region in the D-Spec genome. The amplified fragment was used for the CRISPR/Cas9 with gRNA plasmid in *E. coli* DH5α cells with *cscBKA* operon inserted. The deletion was confirmed by colony PCR with primers del_flagel_F/Del_flagel_R. The gel for confirmation of this deletion is represented in Fig. S3.

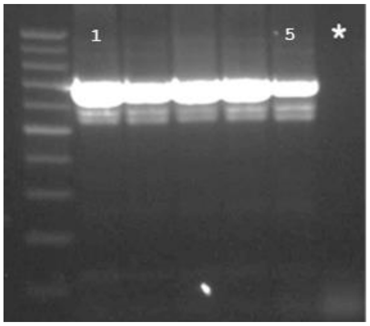


**Fig. S3.** Colony PCR for confirmation of deletion of 40.4 kb. Primers: flagel_seq_F/flagel_seq_R. (*) represents blank.The numbers (1-5) indicate the colonies tested. 100 bp ladder.

#### Cloning in *Saccharomyces cerevisiae*

For yeast plasmid cloning for donor amplification, pEasyG3-mic and pEasyG3-nat or pEasyG4-hph were used as backbone for cloning. The 2-micron origin of replication was amplified by PCR using 2Mic_f/2micMX primers from pEasyG3-mic (1,452 bp), while the resistance marker was amplified with tMX_2m/MXProUni primers from pEasyG3-nat or hph (1,449 bp or 1,645 bp).

##### *galK::cscBKA*

The left and right of *galK* flanking regions were amplified by PCR with mx_galK/AmpTerm_R (594 bp) and cscB_galK_F/galK_2mic (506 bp) primers from the genomic DNA of cells containing the two GFP copies of integrated. The *cscBKA* operon was amplified by PCR from the lineage with this operon integrated with primers term_cscA_F/galK_cscB_R (3,880 bp). These reactions were transformed together with the origin of replication and the resistance marker into *S. cereviseae* cells. Colony PCR confirmed plasmid assembly using cscBterm/right_R (318 bp) and galK_amp/cscA_f (285 bp) primers.

The donor was amplified by PCR with galK_left_f/Right_rev (5,087 bp) primers and used in the CRISPR/Cas9 technique. Colony PCR confirmed insertion using cscBterm/galT_rev (540 bp) and galM_f/cscA_f (762 bp) primers. Gel confirmation is shown in Fig. S4.

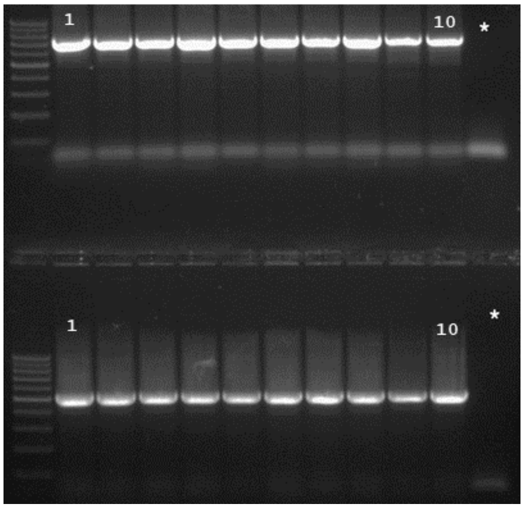


**Fig. S4.** Colony PCR for confirmation of insertion of *cscBKA* operon in the *galK* gene. Primers: galM_f/cscA_f (762 bp, up) and cscBterm/galT_rev (540 bp, down). The numbers (1-10) indicate the colonies tested. (*) represents blank. 100 bp ladder.

##### *cscA::sucP* and *cscA::pJ23119-sucP*

For cloning of pMIC_cscBKsucP, the *sucP* gene with the flanking SS9 region was amplified by PCR from plasmid pUC57_sucP with SS9R_2M/SucP_fw (2,056 bp) primers. The fragment containing *cscBK* genes and the other flanking region was amplified by PCR from pCscBKA, using SS9L_MX/sucP_rev (2,999 bp) primers. For pMIC_cscBKsucP(J23119), primers J23_f/SS9R_2M (2,116 bp) from pUC57_sucP and primers SS9L_MX/CscK_r (3,048 bp) were used. These fragments were transformed into yeast. Colony PCR using CscA_f2/SucP_r (312bp for *cscA::sucP* and 347 bp for cscA::*pJ23119-sucP*) primers, confirmed plasmid assembly. The donor was amplified by PCR with primers csc_F6/SS9Right1 (2,413 bp for *cscA::sucP*; 2,438 bp for *cscA*::*pJ23119-sucP*) and used for CRISPR/Cas9 technique in *E. coli* DH5α cells with *cscBKA* operon inserted. Insertion was confirmed by colony PCR with primers csc_F6/SucP_r (423 bp for pMIC_cscBKsucP(J23119) and 388 bp for pMIC_cscBKsucP) and B0015_f2/FD1 (524 bp). The gel for confirmation of these insertions shown in Fig. S5.

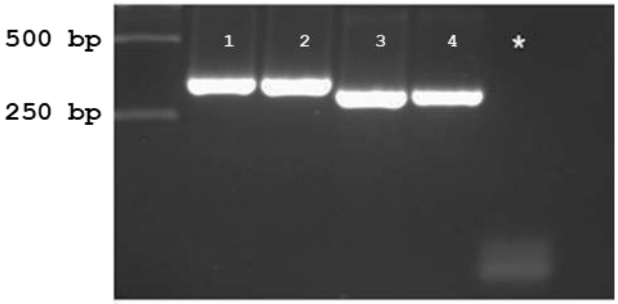


**Fig. S5.** PCR from genomic DNA for confirmation of insertion of *sucP* operon in the *cscA* gene, with prJ23119 (1 and 2) and only with natural promoter (3-4). Primers: CscA_f2/SucP_r (312bp for *cscA::sucP* and 347 bp for cscA::*pJ23119-sucP*) . (*) represents blank.

##### *lacZ::sucP*

For this cloning, the left and right flanking regions were amplified from genomic DNA of cells with two integrated GFP copies using mx_lacZ/Ampterm_R (594 bp) and J23_lacZ_F/lacZ_2mic (442 bp) primers. The sucP gene was amplified from pUC57_sucP with galK_sucP_F/galK_J23_R (1,602 bp) primers. These reactions were transformed together with the origin of replication and the resistance marker into *S. cereviseae* cells. Plasmid assembly was confirmed by PCR with lacZ_amp_f/SucP_R2 (259 bp) and SucP_r/donor_lacZ_r (501 bp) primers. The donor was amplified by PCR with lacZ_left_f/donor_lacZ_r (2,530 bp) primers and used in the CRISPR/Cas9 technique. Insertion was confirmed by PCR with SucP_r/LacZ_R (553 bp) and SucP_R2/LacZ_F (698 bp) primers. Confirmation of the insertion is shown in Fig. S6.

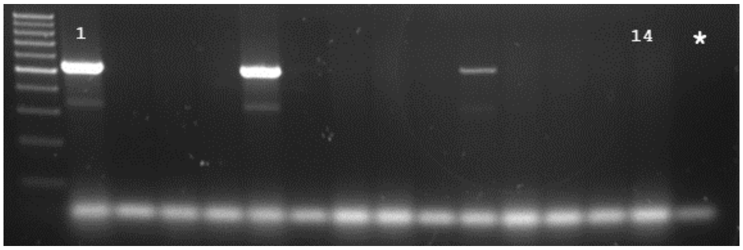


**Fig. S6.** Colony PCR for confirmation of insertion of *sucP* operon in the *lacZ* locus. Primers: SucP_r/LacZ_R (553 bp). The numbers (1-14) indicate the colonies tested. (*) represents blank. 100 bp ladder.

##### *galK*::*gfp*

The mEGFP coding sequence was amplified by PCR from the mEGFP_pBAD plasmid with the primers termAmp_GFP_f/RBS_GFP_F (772 bp). The right and left flanking regions, with the J23119 promoter and AmpR terminator, respectively, were amplified by pMIC_AmpR_ΔgalK, with mx_galK/Ampterm_R (594 bp) and RBS_J23_R/galK_2mic (486 bp) primers. These fragments were transformed into *S. cerevisiae*, and positive cloning were selected by colony PCR with galK_amp/GFP_f (307 bp) and GFP_r/donor_lacZ_r (548 bp) primers. After growth and purification, the donor was amplified by PCR using lacZ_left_f/Right_rev (1,757 bp) primers, which was co-transformed with gRNA plasmid into *E. coli* DH5α with the *cscBKA* operon integrated into the genome and Cas9 plasmid. Insertions confirmation was performed by colony PCR using primers galM_f/GFP_f (783 bp) and GFP_r/galT_rev (608 bp) primers. Confirmation of the insertion is shown in Fig. S7.

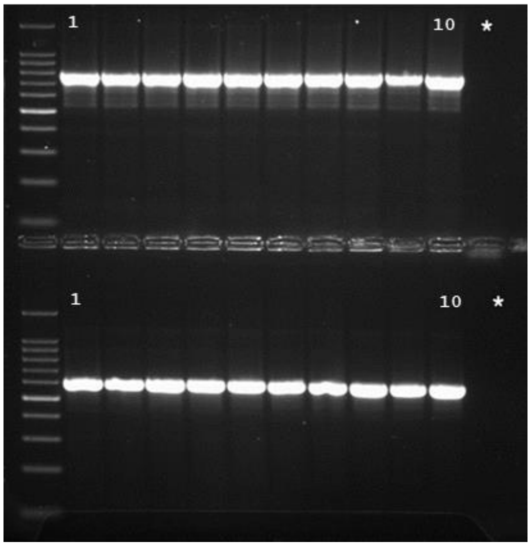


**Fig. S7.** Colony PCR for confirmation of insertion of *gfp* in the *galK* locus. Primers: galM_f/GFP_f (783 bp, up) and GFP_r/galT_rev (608 bp, down). The numbers (1-10) indicate the colonies tested. (*) represents blank. 100 bp ladder.

##### *lacZ::gfp*

The mEGFP coding sequence was amplified by PCR from the mEGFP_pBAD plasmid using termAmp_GFP_f/RBS_GFP_F (772 bp) primers. The right and left flanking regions, containing the J23119 promoter and AmpR terminator, respectively, were amplified by from pMIC_AmpR_lacZ, with mx_lacZ/Amp_term_R (594 bp) and RBS_J23_R/lacZ_2mic (486 bp) primers. These fragments were transformed into *S. cerevisiae*, and positive cloning was selected by colony PCR with lacZ_amp_f/GFP_f (307 bp) and GFP_r/donor_lacZ_r (548 bp) primers. After growth and purification, the donor was amplified by PCR using lacZ_left_f/donor_lacZ_r (1,757 bp) primers and co-transformed with gRNA plasmid into *E. coli* DH5α with integrated *cscBKA* operon and Cas9 plasmid. Insertions confirmation was performed made by colony PCR using LacZ_F/GFP_f (746 bp) and GFP_r/galT_rev (600 bp) primers. Confirmation of the insertion is shown in Fig. S8.

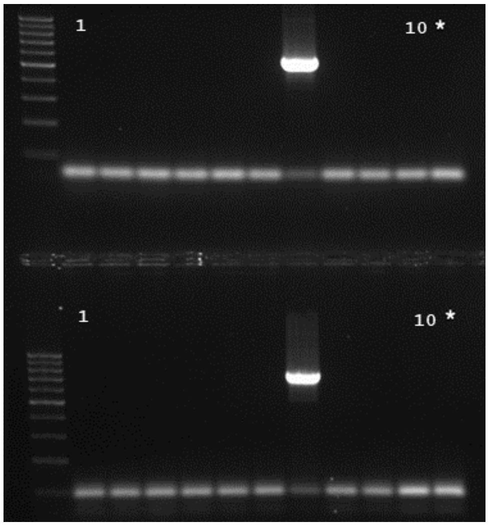


**Fig. S8.** Colony PCR for confirmation of insertion of *gfp* in the *lacZ* locus. Primers: GFP_r/LacZ_R (600 bp, up) LacZ_F/GFP_f (746 bp, down). The numbers (1-10) indicate the colonies tested. (*) represents blank. 100 bp ladder.

##### *lacZ::phaCAB*

For this cloning, the left and right flanking regions were amplified by PCR using mx_lacZ/Amp_term_R (594 bp) and phaC_RBS_F/lacZ_2mic (506 bp) primers from genomic DNA of cells with two copies of GFP integrated into the genome. The *phaCAB* operon was amplified from the pBBRMCS-2::phaCAB plasmid with term_phaB_F/RBS_phaC_R (3,880 bp) primers. These reactions were transformed together with the origin of replication and the resistance marker into *S. cereviseae* cells. Plasmid assembly was confirmed by colony PCR using lacZ_amp_f/phaB_rev (254 bp) and phaC_fwd/donor_lacZ_r (529 bp) primers. The donor was amplified by PCR with the lacZ_left_f/donor_lacZ_r (4,888 bp) primers and used in the CRISPR/Cas9 technique. The insertion was confirmed with primer pairs phaC_fwd/LacZ_R (581 bp) and phaB_rev/LacZ_F (698 bp). Confirmation of this insertion is shown in Fig. S9.

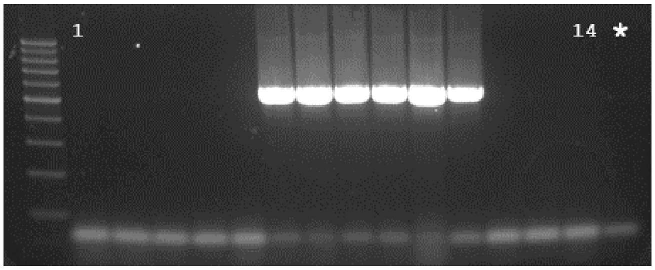

**Fig. S9.** Colony PCR for confirmation of insertion of *phaCAB operon* in the *lacZ* locus. Primers: phaC_fwd/LacZ_R (581 bp). The numbers (1-14) indicate the colonies tested. (*) represents blank. 100 bp ladder.

***galK*::*ampR***

The ampicillin resistance marker was amplified by PCR from the plasmid pTarget_Amp with galK_amp/galK_J23_R (1,103 bp) primers, with the promoter (J23119), the coding region and the terminator. The left and right flanking regions were amplified from the genomic DNA of *E. coli* DH5ɑ of *galK* region, with primer pairs mx_galK/Amp_galK_R (439 bp) and J23_galK_F/galK_2mic (442 bp), respectively. The obtained fragments were mixed with the backbone fragments and used to transform competent *S. cereviseae* cells. Colony PCR confirmed correct cloning with Left_f/AmpT_f (364 bp) and AmpMet/right_R (306 bp) primers. The donor was amplified by PCR with galK_left_f/Right_rev (1,866 bp) primers, purified and used to transform with gRNA plasmid in competent cells of *E. coli* DH5α with *cscBKA* operon, expressing Cas9. Colony PCR was used to demonstrate positive insersion with galM_f/AmpT_f (530 bp) and AmpMet/galT_rev(602 bp) primers. Confirmation of this insertion is shown in Fig. S8.

***lacZ*::*ampR***

J23119 promoter, coding region, and terminator, was amplified by PCR from the pTarget_Amp plasmid using lacZ_amp_f/lacZ_J23_R (1,103 bp) primers. The left and right flanking regions of the *lacZ* locus were amplified from *E. coli* DH5ɑ genomic DNA using mx_lacZ/Amp_lacZ_R (439 bp) and J23_lacZ_F/lacZ_2mic (442 bp) primers. The fragments were mixed with the backbone fragments and used to transform competent *S. cereviseae* cells. Colony PCR was used to confirm correct cloning with lacZ_left_f/AmpT_f (437 bp) and AmpMet/donor_lacZ_r (468 bp) primers. The donor was amplified by PCR with lacZ_left_f/donor_lacZ_r (1,866 bp) primers, purified, and co-transformed with gRNA plasmid into *E. coli* DH5ɑ cells with *cscCBKA* integrated containing the Cas9 plasmid. Colony PCR confirmed positive insertion using LacZ_F/AmpT_f (494 bp) and AmpMet/LacZ_R (520 bp) primers. Confirmation of this insertion is shown in Fig. S10.

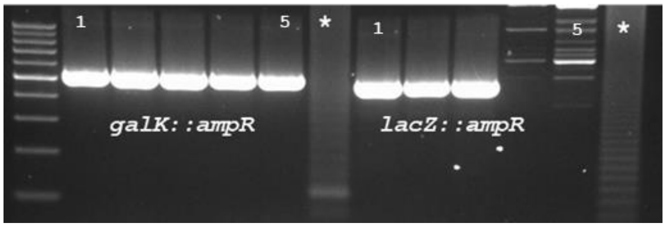


**Fig. S10.** Colony PCR for confirmation of insertion of *ampR* in the *lacZ* and *galK*. Primers for *galK*: galM_f/AmpT_f (530 bp); for *lacZ* LacZ_F/AmpT_f (494 bp). The numbers (1-5) indicate the colonies tested. (*) represents blank. 100 bp ladder.

##### *ΔfliA* (205 bp)

The left and right flanking regions were amplified by PCR from *E. coli* DH5α genomic DNA using mx_fliA/FliA_R (380 bp) and FliA_F/mic_fliA (348 bp) primers, respectively. These fragnents were mixed with the origin of replication and the resistance marker and transformed into *S. cereviseae* cells. Colony PCR using Del_fliA_F/del_fliA_R (279 bp) primers was used to confirm correct cloning. The donor fragment was amplified using Donor_FliA_F/Donor_FliA_R (634 bp) primers and co-transformed with gRNA plasmid into *E. coli* DH5α with the *cscBKA* operon integrated and Cas9 plasmid. Genomic modification was verified by PCR using primers Del_fliA_F/Check_FliA_R (474 bp for deletion, 679 bp for WT). Confirmation of this deletion is shown in Fig. S11.

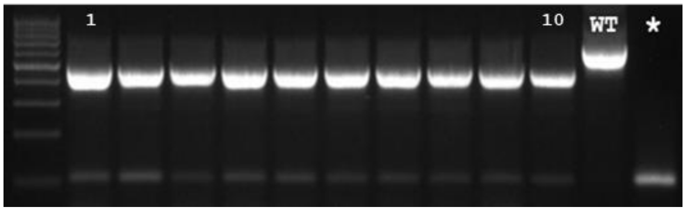


**Fig. S11.** Colony PCR for confirmation of deletion of 205 bp on *fliA*. Primers: Del_fliA_F/Check_FliA_R (deletion 474 bp; WT: 679 bp). The numbers (1-10) indicate the colonies tested.(*) represents blank; WT: wild type. 100 bp ladder.

##### *nudJ (G115V)*

To clone the SNP region of the *nudJ* gene, the flanking regions were amplified by PCR using the primers pairs mx_nudJ/nudJ_R (443 bp) and nudJ_F/mic_nudJ_R (452 bp). Mutation, restriction site, and PAM decharacterization was introduced by homologous primers nudJ_F and nudJ_R. These products were transformed with the origin of replication and the resistance marker into *S. cereviseae* cells. PCR confirmation of the plasmid assembly was done using nudJ_seq_F/nudJ_seq_R (309 bp) primers. The donor was amplified by PCR with Donor_nudJ_F/donor_nudJ_R (814 bp) primers and co-transformed with gRNA plasmid into *E. coli* DH5α with the *cscBKA* operon integrated and Cas9-expressing plasmid. Selection of bacterial colonies with genomic modification was performed by colony PCR using primers XbaI_SNP_nudJ/nudJ_XbaI (269 bp). After confirming amplification, fragments were digested with XbaI. The positive modification was confirmed by forming a 210 bp fragment. The gel confirming this single nucleotide change is shown in Fig. S12.

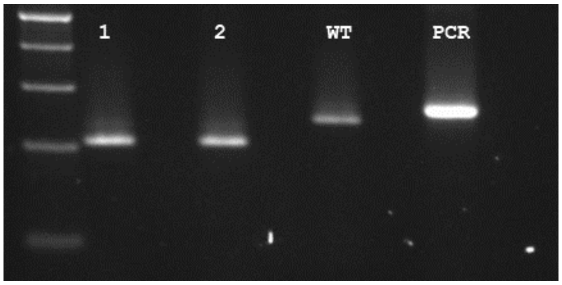


**Fig. S12.** Digestion of fragment amplified by PCR from genomic DNA for confirmation of nucleotide change in *nudJ* gene. The positive modification was confirmed forming a 210 bp fragment. The numbers (1-2) indicate the clones tested. Wd: without digestion; WT: wild type digested. 100 bp ladder.

##### ***purB* (L115G)**

To clone the region with the SNP of the *purB* gene, the flanking regions were amplified by PCR with primers pairs mx_purB/purB_R (375 bp) and 2purB_F2/mic_purB_R (409 bp). These reactions were co-transformed with the origin of replication and the resistance mark into *S. cereviseae* cells. Mutation and PAM decharacterization was specified by homologous primers 2purB_F2 and purB_R. Mutation in the purB gene naturally generated a restriction site for EcoRI. Colony PCR with primers purB_seq_F/purB_seq_R (263 bp) confirmed cloning.  The donor was amplified by PCR with donor_purB_F/Donor_purB_R (695 bp) primers, and was co-transformed with gRNA plasmid into *E. coli* DH5α with the *cscBKA* operon integrated and Cas9-expressing plasmid. Selection of bacterial colonies with genomic modification was performed with colony PCR with purB_seq_F/Ecori_purb (325 bp) primers. Corrected amplifiied fragments were digested with EcoRI. The positive modification was confirmed by forming a 179 bp fragment. The gel confirming this single nucleotide change is shown in Fig. S13.

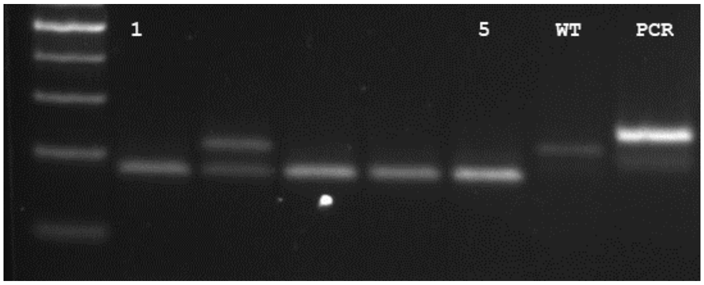


**Fig. S13.** Digestion of fragment amplified by colony PCR for confirmation of nucleotide change in *purB* gene. The positive modification was confirmed, forming a 179 bp fragment. The numbers (1-5) indicate the colonies tested. Wd: without digestion; WT: wild type digested. 100 bp ladder.

#### Modifications using the EasyOligo approach

##### *ΔflhDC*

The oligo-donor was amplified by PCR with primers L-FlhC/mid-FlhCD/R-FlhD (160 bp). This PCR product was used in CRISPR/Cas9 with corresponding gRNA plasmid in *E. coli* DH5α cells with the *cscBKA* operon integrated. The confirmation of deletion was made using primers Check_FlhCD_F/Check_FlhCD_R (deletion: 222 bp; wild type: 1,103 bp) in colony PCR. The gel for confirmation of this deletion is represented in Fig. S14.

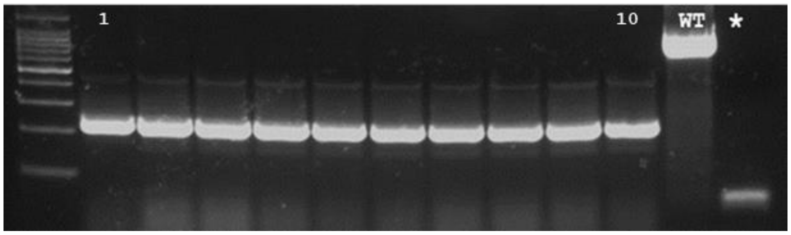


**Fig. S14.** Colony PCR for confirmation of deletion of *flhCD*. Primers: Check_FlhCD_F/Check_FlhCD_R (deletion: 222 bp; wild type: 1,103 bp). The numbers (1-10) indicate the colonies tested. (*) represents blank; WT: wild type. 100 bp ladder.

##### *ΔhflK*

The oligo-donor was amplified by PCR with primers L-hflK/Mid-hflK/R-hflK (160 bp) and employed in CRISPR/Cas9 with the corresponding gRNA plasmid in *E. coli* DH5α cells containing the *cscBKA* operon. Deletion was confirmed by colony PCR using primers Check_hflK_F/Check_hflK_R (deletion: 215 bp; wild type: 1,374 bp). Confirmation of this deletion is shown in Fig. S15.

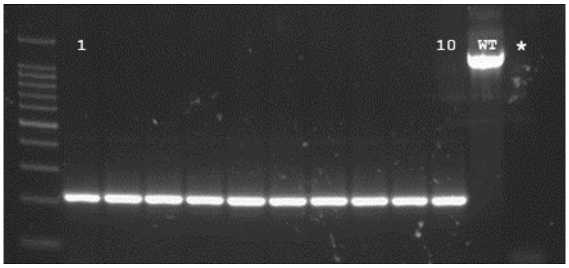


**Fig. S15.** Colony PCR for confirmation of deletion of *hflK*. Primers: Check_hflK_F/Check_hflK_R (deletion: 215 bp; wild type: 1,374 bp). The numbers (1-10) indicate the colonies tested. (*) represents blank; WT: wild type. 100 bp ladder.

**ΔCA (12 kb)**

The oligo-donor was amplified by PCR using primers L-galF/Mid-colonic/R-wcaI (160 bp). and used in CRISPR/Cas9 with the corresponding gRNA plasmid in *E. coli* DH5α cells containing the *cscBKA* operon. Deletion was confirmed Confirmation of this deletion is shown in Fig. S14 colony PCR using primers ChecK_GalK_F/ChecK_GalK_R (deletion: 239 bp). The gel for confirmation of this deletion is represented in Fig. S16.

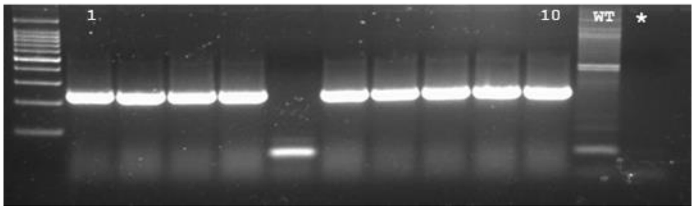


**Fig. S16.** Colony PCR for confirmation of deletion of CA. Primers: ChecK_GalK_F/ChecK_GalK_R (deletion: 239 bp; wild type: ~12 kb). The numbers (1-10) indicate the colonies tested. (*) represents blank; WT: wild type. 100 bp ladder.

***ΔgalK***

The oligo-donor was amplified by PCR with primers galK_L/Oligo central galK/galK_R (140 bp) and used in CRISPR/Cas9 with corresponding gRNA plasmid in *E. coli* DH5α cells with the *cscBKA* operon integrated. Deletion was confirmed by colony PCR using primers CheckK_GalK_F/ChecK_GalK_R (deletion: 246 bp; wild type: 278 bp). The gel confirming this deletion is shown in Fig. 5.

##### *dnaK (T417N)*

The oligo-donor was amplified by PCR with primers L2-dnaK/Mid2-dnaK/R2-dnaK (157 bp). and applied in CRISPR/Cas9 with the corresponding gRNA plasmid in *E. coli* DH5α cells with the *cscBKA* integrated. Nucleotide change was made by digestion with PvuII of fragment amplified by colony PCR using primers DnaK2_PvuII_F/DnaK2_seq_R (264 bp). Positive mutants were identified by the presence of a 159 bp fragment after digestion. Sanger sequencing of positive transformants was performed using PCR amplification with primers DnaK2_seq_F/DnaK2_seq_R (268 bp) in both directions. The gel confirming this single nucleotide change is shown in Fig. S17.

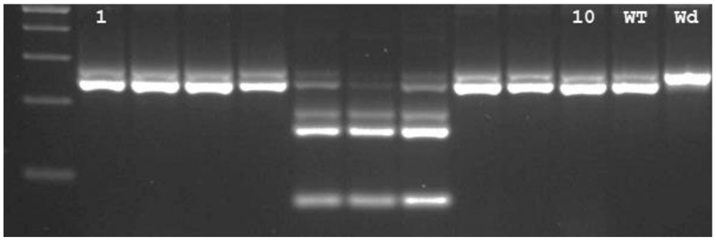


**Fig. S17.** Digestion of fragment amplified by colony PCR for confirmation of nucleotide change in *dnaK* gene. The positive modification was confirmed forming a 159 bp fragment. The numbers (1-10) indicate the colonies tested. Wd: without digestion; WT: wild type digested. 100 bp ladder.

##### *zwf (D405E)*

The oligo-donor was amplified by PCR with primers L-zwf/Mid-zwf/R-zwf (155 bp). This PCR product was used in CRISPR/Cas9 with respective gRNA plasmid in   cellswith *cscBKA* operon integrated. Nucleotide change was confirmed by digestion with PstI of fragment amplified by colony PCR using primers zwf_PstI_F/zwf_seq_R (302 bp). Positive mutants were selected by 174 bp and 102 fragments presence. Sanger sequencing of positive transformants was made using amplification by PCR with zwf_seq_F/zwf_seq_R (218 bp) primers, in both directions. The gel for this modification is presented on Fig. 5.
