## Supplementary Methods for "Engineering Adaptive Alleles for *Escherichia coli* Growth on Sucrose Using the EasyGuide CRISPR System"

**Minimum medium composition**

**5X Minimal medium salts**

| Compound | Concentration |
|---|---|
| $NaH_2PO_4.H_2O$ | 36.4 g/L |
| $KH_2PO_4$ | 15 g/L |
| NaCl | 2.5 g/L |
| $NH_4Cl$ | 5 g/L |

**10,000x Mineral Solution**

| Compound | Concentration |
|---|---|
| $H_3BO_3$ | 5 g/L |
| $MnSO_4$ | 4 g/L |
| $ZnSO_4 \cdot 7H_2O$ | 4 g/L |
| $FeCl_2.6H_2O$ | 2 g/L |
| Molybdic acid | 0,4 g/L |
| KI | 1 g/L |
| $CuSO_4 \cdot 5H_2O$ | 0.4 g/L |
| Citric acid | 10 g/L |

**Quantitative methods**

**Quantification of sucrose consumption**

For sucrose quantification, cells were pre-adapted in minimal medium with sucrose as the sole carbon source at 37 ºC and 100 rpm, in 20 mL using 125 mL shake flasks. The experiment was performed in triplicate in the same conditions as pre-adaptation. After 24 h the samples were collected and frozen Samples were diluted, filtered with a 0.22 μm syringe filter, and quantified using a protocol adapted from de Freitas and Brienzo (de Freitas, C., Brienzo, M., 2023. BioEnergy Research. 16, 1040-1050). XOS concentration was determined by high-performance liquid chromatography (HPLC) using Waters 2414 equipment and an Aminex HPX-87C BIO-RAD column (300 × 7.8 mm) at 80 °C. Ultrapure water with a flow rate of 0.6 mL/min served as the eluent. For quantification, 20 μL of sample was used, and detection was performed by refractive index

**PHB quantification**

After strain cultivation, the culture was centrifuged at 8,000 x g for 20 min at 4 °C and the supernatant discarded. The recovered biomass was lyophilized, and 10 mg of the lyophilized cells were subjected to methanolysis. 2 mL of chloroform, 2 mL of methanol acidified with 15% sulfuric acid (85:15 v/v), and 100 μL of benzoic acid in methanol, with a ratio of 0.02 g/mL, were added to glass test tubes. The mixture was vortexed at 2,500 rpm for 1 min after each addition. The tubes were then incubated in a water bath at 85 °C for 4 hours, with constant shaking after the first hour. After the methanolysis, the tubes were cooled to room temperature, and 2 mL of ultrapure water was added, vortexed for 30 seconds, and incubated for 4 h to separate the organic and aqueous phases. The aqueous phase was discarded, and the organic phase, containing the methyl esters, was stored at -20 °C for later analysis in a GCMS-QP2010 Ultra Mass Spectrometer (Shimadzu), equipped with an Rtx®-5MS column, using benzoic acid as the internal standard.

GCMS analysis was performed using injector temperature at 250 °C, and starting application on the column at 100 °C for 3 min, with an initial column temperature of 100 °C held for 3 min, followed by an 8 °C/min increase until reaching 210 °C, which was maintained for 15 min. The detector temperature was 280 °C, with a constant helium carrier gas flow of 0.8 mL/min. Benzoic acid was used as the internal standard, while PHB and PHBV served as external standards (Sigma codes: 36.350-2 and 40.311-3, respectively).

To quantify the percentage of dry mass of PHB produced by each strain, the results of the areas obtained in the GC-MS were subjected to 3 formulas:

Mass of PHB (g) (M):

$$\frac{A \times 5 \times 10^{-9}}{B} \times 40{,}000$$

Where A represents the area value of PHB and B represents the area of benzoic acid.

PHB mass adjusting with the yield of internal standards (R):

$$\frac{M \times 100}{21.5}$$

Where M represents the value of the PHB mass (g).

PHB produced (%) in relation to dry cell mass (DCM) (P):

$$\frac{R}{0.01} \times 100$$

Where R represents the mass of PHB adjusted with the yield of internal standards.

After quantifying the production of PHB by each strain, the results were analyzed in order to compare the quantity produced by the genetically modified *E. coli* DH5α with *cscBKA* an *phacab* operons inserted and the naturally PHB-producing strain.

To production of PHB films, the lyophilized were dissolved in chloroform, at a ratio of 1 g/20 mL for 48 hours at 60 °C with 200 rpm agitation. Cell debris was removed by ultrafiltration through a 0.2 μm PTFE filter, and the resulting liquid was deposited onto a glass surface to form the film.
